# Amyloid β instigates cardiac neurotrophic signaling impairment, driving Alzheimer’s associated heart disease

**DOI:** 10.1101/2023.07.11.548558

**Authors:** Andrea Elia, Rebecca Parodi-Rullan, Rafael Vazquez-Torres, Ashley Carey, Sabzali Javadov, Silvia Fossati

## Abstract

While a link between cardiovascular risk factors and increased Alzheimer’s disease (AD) risk has been reported, it remains unclear whether AD pathology has a direct effect on cardiac function and myocardial innervation. AD and amyloidosis are known to impair neuronal function and affect brain neurotrophic factors (NGF and BDNF) expression. Amyloid aggregates and neuro-signaling impairments may also expose AD patients to peripheral nervous system deficits, promoting cardiac disorders. Here, we characterize cardiac physiology, amyloid pathology, neurotrophic factors loss, and the impoverishment of cardiac neuronal fibers in Tg2576-AD mice hearts, human cardiomyocytes in culture, and human AD post-mortem left ventricular (LV) heart tissue. We reveal that Tg2576 animals exhibit increased myocardial fibrosis, amyloid β (Aβ) deposition, and brain/heart-axis neurotrophic deficiencies, resulting in myocardial denervation and cardiac dysfunction. Aβ oligomers reduce BDNF expression in both human immortalized and iPSC-derived cardiomyocytes, by disrupting TrkB/CREB signaling. Analysis of human LV AD post-mortem tissue confirmed cell and animal results. Our findings elucidate a previously unknown mechanism of Aβ-induced cardiac neurotrophic signaling dysregulation, underscoring the relevance of heart degeneration in AD.

**Translational Perspective:** This research identified cardiac amyloid pathology, neurotrophic factor depletion, and reduced myocardial nerve function in a transgenic model of cerebral amyloidosis (Tg2576) and in human AD heart tissue. These findings carry significant diagnostic and therapeutic implications, emphasizing the role of neuro-signaling disruption in cardiac physiology impairment linked to AD. Our study advocates for considering cardiac complications in AD management and paves the way for future precision medicine approaches to enhance systemic clinical strategies for treating AD, proposing the cardiac neurotrophic signaling pathway as a potential therapeutic target.

## 1. Introduction

Alzheimer’s disease (AD) is an escalating global health concern, anticipated to triplicate in prevalence by 2050, primarily due to population aging^1^. This complex neurodegenerative disorder is characterized by the progressive accumulation of insoluble amyloid-β (Aβ) plaques in the brain parenchyma together with neurofibrillary tangles formed by hyperphosphorylated tau. Besides Aβ plaques and tau buildups, about 90% of AD patients also show Aβ deposits in and around the cerebral vasculature, known as cerebral amyloid angiopathy (CAA). CAA contributes significantly to the progression of AD pathology by promoting neuroinflammation, neurovascular dysfunction, decreased brain clearance, increased blood-brain barrier (BBB) permeability, and eventually neurodegeneration^2,3^.

Despite the main symptoms of the disease predominantly concerning the central nervous system (CNS), emerging evidence highlights extracerebral implications of AD, involving peripheral organs such as the heart^4^. Although abundant attention has been given to the impact of cardiovascular risk factors on AD risk and progression^5–7^, only recent studies suggest a direct effect of AD pathology on cardiac dysfunction, and reveal Aβ peptides (Aβ40 and Aβ42) and big tau accumulation in the cardiac tissue of AD patients^8,9^.

The accumulation of Aβ in the AD brain is known to significantly affect the expression of the two main neurotrophic factors, nerve growth factor (NGF) and brain-derived neurotrophic factor (BDNF) ^10^. This neurosignaling pathway’s maladaptive remodeling is associated with cognitive decline^11^. The production of neurotrophins such as BDNF is also important for peripheral organs and cells, including endothelial cells, muscle cells, and cardiomyocytes. BDNF operates through tropomyosin receptor kinase B (TrkB) stimulation, contributing to the activation of neuronal pro-survival genes, including BDNF itself, through cAMP response element-binding protein (CREB) signaling^12^. Additionally, BDNF exerts pleiotropic effects in endothelial cells, muscle cells, and cardiomyocytes. BDNF/TrkB/CREB signaling is crucial for proper heart development and function in adulthood^13^. It is plausible to hypothesize that the cerebral neurotrophic signaling degeneration in AD may be accompanied by a gradual decline also in peripheral levels and function of these neuromodulators. A lack of neurotrophic support may result in a severe derangement of the cardiac nervous system, potentially culminating in severe heart dysfunction. Notably, declining circulating BDNF levels have also been linked to cardiac disease progression, particularly of ischemic origin^14^.

However, the effects of AD pathology on the heart pathophysiology remain poorly understood, and whether and through what mechanisms AD and amyloidosis modulate NTFs levels and innervation in the cardiac tissue have not been previously explored. Also, whether specific Aβ aggregates such as toxic oligomers, which may accumulate in the AD heart, fuel cardiac neuro-signaling dysregulation, is still to be defined.

To address these questions, we analyzed whether and how the accumulation of cardiac Aβ aggregates affected peripheral innervation and NTFs expression in the heart of Tg2576 transgenic mice, one of the most widely used models of cerebral amyloidosis. We observed a progressive decline in cardiac function, that became significant in 13 months-old Tg mice compared to controls, and was accompanied by Aβ deposition, fibrosis, increase in cardiac mass and reduced contractility. We found a significant reduction in cardiac nerve fiber density and neurotrophins levels, adrenergic and neuronal fibers regeneration markers, in Tg2576 mice compared to controls. Through *in vitro* experiments on human AC16 cardiomyocytes, neuroblastoma cells, and validation in human-induced pluripotent Stem Cell-derived cardiomyocytes (hiPSC-CMs), we showed that exposure to Aβ oligomers significantly reduced the expression of BDNF in both neural and cardiac cells. BDNF acts through tropomyosin receptor kinase B (TrkB) stimulation, contributing to the activation of pro-survival genes, including BDNF itself, through cAMP response element-binding protein (CREB) signaling. We demonstrated that CREB levels were reduced in cardiomyocytes exposed to Aβ oligomers, and that a TrkB agonist could mitigate caspase 3 activation and apoptosis in Aβ-exposed cardiomyocytes. Finally, we validated these findings by analyzing the neurotrophic cardiac profile in human left ventricle (LV) post-mortem tissue from AD patients with CAA compared to age-matched heathy subjects, thus opening novel avenues for future investigations into the impact of AD pathology on cardiac physiology.

This study demonstrates that pathological Aβ oligomers induce cardiac neuro-signaling impairment in mouse models and human AD hearts, elucidates the molecular mechanism responsible for Aβ-mediated cardiac neurotrophic deficiency and nerve fiber loss in AD, and may suggest novel therapeutic avenues to tackle this devastating disease, contributing to filling an important gap on the effects of AD on cardiac function and pathology.

## 2. Methods

### Animal Model

Male and female Tg2576 mice, a model of cerebral amyloidosis bearing the Swedish mutation (KM670/671NL), and age-matched wild-type (WT) littermates were bred in-house. Mice were maintained under controlled conditions (∼22°C and in a 12-hour light-dark cycle, lights from 7 am to 7 pm) with unrestricted access to food and water. The generation of B6; SJL-Tg (APPSWE)2576Kha mice (Tg2576) on a B6; SJL Mixed Background was described in^15^. Tg2576 mice are known to develop numerous cerebral parenchymal amyloid-β plaques at 11-13 months and CAA at 10-11 months^16^. All experiments and animal protocols were performed in equal numbers of male and female mice, according to procedures approved by the Institutional Animal Care and Use Committee of Temple University School of Medicine and conformed to the National Research Council Guide for the Care and Use of Laboratory Animals published by the US National Institutes of Health (2011, eighth edition).

### Assessment of cardiac function by transthoracic echocardiography

Transthoracic bi-dimensional M-mode echocardiographic analysis was accomplished in 4, 8 and 13 months-old mice (before and after the development of brain Aβ pathology), under anesthesia, using a VisualSonics Vevo 2100 system (VisualSonics, Toronto, Canada). Before echocardiographic evaluation, the chest of the mice was carefully depilated by an operator. Subsequently, the mice were positioned in a clinostatic orientation on a temperature-controlled surface, with ECG leads embedded for monitoring, ensuring a constant temperature of 37°C. Anesthesia was then administered to the animals via an anesthetic chamber, utilizing isoflurane (Zoetis IsoFlo, Kalamazoo, MI), with an induction concentration of 3.0% and a maintenance range of 1–2.5%. This anesthesia regimen was meticulously managed to keep the animals within the desired anesthesia depth throughout the procedure, optimizing their physiological stability. Cardiac function and structure were evaluated by measuring left ventricle (LV) wall thickness, end-diastolic diameter (LVEDD), end-systolic diameter (LVESD), end-diastolic volume (LVEDV), end-systolic volume (LVESV), ejection fraction (EF%), and fraction shortening (FS%) percentage^17^. In accordance with established protocols^18^, speckle tracking-based strain echocardiography (STE) was conducted using a parasternal long-axis B-mode loop captured through the VisualSonics Vevo 2100 system (VisualSonics). High-speed imaging was ensured, with all recordings comprising 300 frames at a rate exceeding 200 frames per second, digitally stored as cine loops. Subsequently, the images obtained underwent comprehensive analysis using Vevo Strain Software (Vevo LAB 1.7.1). LV endocardial strain was computed in both radial and longitudinal axes and expressed as peak strain percentage. Additionally, LV deformation rate over time was evaluated and presented as strain rate (SR; peak 1/s). Global and regional assessments of LV endocardial longitudinal and radial strain, along with SR, were performed based on the average measurements derived from six LV segments (comprising basal, mid, and apical anterior segments, as well as basal, mid, and apical posterior segments). Regional LV endocardial longitudinal and radial strain, along with SR, were analyzed individually for each LV segment and then averaged across both the anterior and posterior walls. Vector diagrams illustrating the direction and magnitude of endocardial deformation were acquired from parasternal long-axis B-mode images and subjected to detailed analysis. Diastolic dysfunction was assessed by utilizing the reverse peak option to evaluate reverse global longitudinal strain (GLS) and global radial strain (GRS) rates, which were derived from long-axis B-mode traces^18,19^. Lastly, animals were euthanized at 13 months for organ explant and biomolecular and histological analysis.

### Tissue processing and morphological evaluations

Cardiac specimens were fixed in Zamboni’s solution (2% paraformaldehyde and picric acid) overnight at 4°C, then cryoprotected with 20% sucrose in PBS for 24 hours at 4°C and cut in 5-µm-consecutive thick longitudinal sections using a cryotome (Leica 2000R, Germany). Interstitial fibrosis was assessed via Picrosirius Red Stain Kit (Abcam, ab150681), and images were acquired using a Nikon NiE Fluorescence Microscope with an 20X objective^20^. Cardiac fibrosis density was measured using ImageJ software (NIH, version 1.30) and expressed as a percentage of fibrotic area over the total area. A single operator blindly executed all the quantifications.

### Immunostaining

The cardiac sections were processed via the indirect immunofluorescence method using rabbit polyclonal antibodies against tyrosine hydroxylase (TH, Millipore, AB152; 1:100) to stain adrenergic nerve fibers and growth association protein 43 (GAP-43, Millipore, #AB5220; 1:100) for regenerating nerve terminals. Cardiac nerve fibers showed a regular pattern along cardiomyocytes^21^. Heart nerve ending density was quantified using a square grid with 24 probes (area: 1 cm^2^ each) overlapped to the digital images, which were acquired using a Nikon Eclipse Ti fluorescence microscope with 20X and 60X objectives. Five areas were randomly chosen for quantification from each sample’s three sections. The total number of intercepts between nerve fibers and probes for each area was assessed, and cardiac nerve fiber density was measured and expressed as the mean number of intercepts per area (fibers/mm^2^). A single operator assessed all the measurements blindly. Next, to evaluate amyloid-β accumulation, 5-μm-thick cardiac sections from each experimental group were marked with 6E10 anti-Aβ antibody (Covance, SIG-39320; 1:500) and co-stained with cardiac troponin T (cTnT, Thermo Scientific, #MA5-12960; 1:250), collagen type I (Rockland Immunochemicals, #600-401-103-0.1; 1:100), cardiac Troponin I (cTnI, Abcam, ab47003; 1:200), or cleaved caspase-3 (Asp175) (Cl-Casp-3, Cell Signaling, #94530; 1:200) antibodies. Additionally, to characterize βsheet (fibrillar) amyloid deposits, cardiac sections were stained with Thioflavin-S solution (Acros organics, 213150250; 0.5g of Thioflavin-S powder in 100 ml dH₂O) for 30 minutes at room temperature shielding from the light. Next, the sections were dehydrated in 80% ethanol and thus washed in PBS 1X buffer. Nuclei were labeled with 4’, 6-diamidino-2-phenylindole (DAPI, Sigma Aldrich, D9542; 1:5000), and sections were mounted using an aqueous mounting medium. Digital images were captured and analyzed by a Nikon Eclipse Ti fluorescence microscope with a 60X objective. To evaluate vascular components, after fixation, cardiac specimens were cut into 5-µm-thick longitudinal sections using a cryotome (Leica 2000R, Germany). Next, tissue sections were incubated for 30’ minutes at 4°C in a blocking solution (10% donkey serum supplemented with 0.3% Triton X-100 in phosphate-buffered saline). Via indirect immunofluorescence procedure, sections were analyzed using goat polyclonal antibodies against CD31 (R&D Systems, AF3628-; 1:30) and α-smooth muscle actin (α-SMA, Abcam, ab32575; 1:500) at 4°C overnight, followed by specific secondary antibodies conjugated with cyanine 3 or cyanin 5 respectively, to stain vascular network. For serum amyloid A3 immunostaining (SAA3), fixed 5-µm-thick cardiac sections were retrieved with formic acid 70% solution (Thermo Scientific, #270480010) for twenty minutes at room temperature, they were then stained using mouse monoclonal antibody against SAA (Abbexa, abx018140, 1:50) for two hours at 37°C, followed by specific secondary antibodies combined with Alexa Fluor 488-conjugated anti-mouse IgG. DAPI was used to counterstain nuclei and cardiac sections were mounted with a Vectashield vibrance antifade mounting medium (H-1000). To assess the changes in the cross-sectional area (CSA) of cardiomyocytes, 5μm-thick cardiac sections from each experimental group were incubated for 30 minutes at 4°C in a blocking solution (10% donkey serum) and permeabilized with 0.3% Triton X-100 in phosphate-buffered saline. The tissue sections were then stained with 5 μg/mL wheat germ agglutinin (WGA, #W11261, Thermo Fisher Scientific, Waltham, MA, USA) as previously described^22^ (PMID: 34404991). Afterward, the sections were mounted with Vectashield Vibrance antifade mounting medium (#H-1000). Digital images were captured and analyzed using a Nikon Eclipse Ti fluorescence microscope with a 20X objective. Five to six randomly selected fields per sample were analyzed, totaling 50-60 cardiomyocytes per section (10 cells per area). The mean cardiomyocyte diameter (μm) was measured using the ImageJ software. Lastly, to assess amyloid-β deposits in the human heart, 5-μm-thick LV sections from healthy donors and AD with CAA patients were stained with 6E10 anti-Aβ antibody (Covance, SIG-39320; 1:500) and co-stained with cardiac Troponin I (cTnI, Abcam, ab47003; 1:200). Images were acquired and examined using a Nikon Eclipse Ti fluorescence microscope with 20X and 60X objectives. A single operator blindly performed all the procedures.

### Preparation of cell and tissue lysates and immunoblotting analysis

Cellular and cardiac samples were homogenized using RIPA cell lysis buffer (Thermo Scientific) enriched with phosphatases and protease inhibitors (Thermo Scientific). Then, the lysates underwent sonication and were centrifuged for 10 minutes at 4°C at 13,000 rpm to discard the insoluble debris. Next, total protein amounts were quantified via a dye-binding Pierce BCA protein assay kit (Thermo Scientific) and detected using a spectrophotometer reader (SpectraMax i3x Multi-mode Microplate Reader, Molecular Devices) at a wavelength of 562 nm. Equal yields of protein (20-40 μg) were separated through SDS-PAGE and identified by western blot (WB) analysis. Total lysates were used to evaluate the protein levels of NGF (Alomone labs, AN-240; 1:1000), BDNF (Alomone labs, ANT-010; 1:1000), GAP-43 (Millipore, #AB552; 1:1000), Cleaved Caspase-3-Asp175 (Cl-Casp-3, Cell Signaling, #94530; 1:500), dopamine β hydroxylase (DβH, Millipore, #AB1536; 1:1000), human Aβ [mouse monoclonal anti-Aβ antibodies mixture composed of 4G8 epitope (residues Aβ18-22, Covance, SIG-39320; 1:1000) and 6E10 epitope (residues Aβ3-8, Covance, SIG-39220; 1:1000)], serum amyloid A3 (SAA3, Abcam, #ab231680; 1:500) CREB (Cell Signaling, #9197; 1:1000), fibronectin (Abcam, #ab2413; 1:1000), matrix metalloproteinases-2 (MMP-2, Cell Signaling, #35814; 1:1000), Laminin A/C (Santa Cruz Biotechnology, sc-376248; 1:500), Actin (Millipore, MAB1501; 1:1000) and GAPDH (Santa Cruz Biotechnology, sc-32233,6C5; 1: 2000), the three latter of which were used as loading controls. Protein bands were detected using Odyssey® CLx Imaging System according to the manufacturer’s instructions and quantified with Image Studio™ Lite Software^23^.

### ELISA

Aβ40 and Aβ42 peptide levels were detected using a commercial ELISA kit (Invitrogen KHB3481 and KHB3441, respectively). Briefly, cardiac specimens for each mouse were collected at the end of the experimental study and homogenized with RIPA cell lysis buffer (Thermo Scientific), supplemented with phosphatases and protease inhibitors (Thermo Scientific, Waltham, MA). Then, ELISA was performed using an equal amount of total cardiac protein following the company’s specifications, and the plate was evaluated at 450 nm with a spectrophotometer reader (SpectraMax i3x Multi-mode Microplate Reader, Molecular Devices).

### Aβ peptides

Pre-aggregated Aβ40 and Aβ42 oligomers were generated from HFIP-pretreated and lyophilized peptides as previously described^24^. Briefly, oligomer enriched Aβ40 and Aβ42 preparations were obtained after a dilution of the lyophilized peptide in DMSO at a concentration of 5 mM, then diluted to 100 μM in ice-cold DMEM/F12 media and incubated at 4°C for 16 hours. Next, Aβ40 and Aβ42 oligomers were resuspended in DMEM/F12 medium to 10 μM as a final concentration for the cell treatments.

### Cell culture and treatments

Human ventricular myocardial cardiomyocytes (AC16, purchased from Millipore, #SCC109) were expanded in 10 cm^2^ dishes not further than the 10th passage, in a humidified environment at 37°C with 5% CO_2_ in DMEM/F12 medium (Lonza Ltd. Basel, Switzerland) supplemented with 12.5% fetal bovine serum (FBS) and 1% penicillin-streptomycin (P/S, Lonza). AC16 cells were grown in the pre-contractile developmental stage, thus allowing us to better track molecular events and cellular responses^25,26^, without the complications and variability associated with isolated adult cardiomyocyte cultures. Human SH-SY5Y neuroblastoma cells (ATCC, CRL-2266, a neuronal cell line used widely in experimental neurological studies^27–29^) were grown in 10 cm^2^ dishes within the 10th passage in a humidified environment at 37°C with 5% CO_2_ in DMEM/F12 medium (Lonza Ltd. Basel, Switzerland) supplemented with 10% fetal bovine serum (FBS) and 1% penicillin-streptomycin (P/S, Lonza). For both cell types, AC16 and SH-SY5Y, an equal number of cells were seeded in 6 well plates, and after 24 hours, once they reached about 70-80% confluency, cells were challenged with human Aβ40 or Aβ42 oligomers (10 μM) for 16 hours. In certain studies, cells were treated with human Aβ40 oligomers (10 μM) for either 15 minutes or 1 hour. For other experiments, cells were exposed to the BDNF/TrkB receptor agonist LM22A-4 (100 nM) or the antagonist ANA-12 (30 μM) for 16 hours. LM22A-4 (4607) and ANA-12 (SML0209) were sourced from Tocris Bioscience and Sigma Aldrich, respectively.

### Maintenance and treatment of Human-Induced Pluripotent Stem Cell-Derived Cardiomyocytes (hiPSC-CMs)

A commercially available hiPSC-CM line, iCell Cardiomyocytes^2^ (Catalog number R1220, Kit/lot 12,012; FUJIFILM Irvine Scientific, Inc., United States), was utilized for this study. The cells were seeded into a 0.1% gelatin-coated 12-well plate at a density optimized to form a monolayer (≈210K cells per well) and cultured in a stage incubator (37°C, 5% CO2) following the provider’s guidelines. Experiments were conducted 5–7 days post-plating, ensuring the cells formed a viable, beating monolayer, in accordance with the manufacturer’s instructions (Supplementary Video). Cells from this specific lot were from the same donor (healthy, female, Caucasian, age <18), as verified by FUJIFILM Cellular Dynamics. For the experiment, hiPSC-CMs were treated with human Aβ40 oligomers (10 μM) for 20 hours, after which the cells were collected for biochemical analysis. For imaging, cells were fixed with 4% paraformaldehyde (PFA) for 10 minutes, washed with 1X PBS, and then blocked and permeabilized for 30 minutes using PBS containing 0.1% Triton X-100 and 2% normal donkey serum. hiPSC-CMs were subsequently labeled with a BDNF antibody (Abcam, ab226843; 1:250) and co-stained for cardiac troponin T (cTnT, Thermo Scientific, #MA5-12960; 1:250), with nuclei labeled using DAPI (Sigma Aldrich, D9542; 1:5000). Imaging was conducted using a Nikon Eclipse Ti fluorescence microscope with a 100X objective, with all procedures performed by a single operator in a blinded manner.

### Nuclear protein isolation

AC16 human cardiomyocytes were cultured in 6 well plates until they reached confluence and treated with human Aβ40 oligomers for 15 minutes. To extract the nuclear fractions, cells were collected using buffer A, containing 10 mmol/L HEPES at pH 7.9, 10mmol/L KCl, 0.1mmol/L EDTA, integrated with 25% NP-40X-100 non-ionic surfactant detergent solution, enriched with protease inhibitors. The cell lysate was transferred in a 1.5 ml tube, placed in ice, and centrifuged for 10 min at 4°C at 13,000 rpm to discard the insoluble debris. The supernatant was transferred to a new 1.5 ml tube to preserve the cytosolic fraction, while the pellet was resuspended in fresh prepared buffer B composed of 20mmol/L HEPES at pH 7.9; 0.4mol/L NaCl; 1mmol/L EDTA; 10% glycerol; supplemented with protease inhibitor. Next, the tube containing the nuclei pellet dissolved in buffer B was placed on a rotator for 2 hours at 4°C and then centrifuged at 13,000 rpm for 5 minutes at 4°C. Lastly, the supernatant (nuclear fraction) was transferred to a new 1.5 ml tube and stored at -80°C, ready for the following protein quantification and immunoblot analysis^23^.

### Human LV specimens

Frozen human LV post-mortem tissue from healthy donors and patients with clinical diagnoses of AD and CAA were obtained from the Banner Sun Health Research Institute clinical biobank to investigate the impact of AD pathology on myocardial innervation. The selection criteria of the study population excluded individuals with comorbidities known to substantially affect cardiac function, including high blood pressure, cerebrovascular accidents (e.g., strokes and lacunes), brain/cardiac neoplastic conditions, secondary amyloidosis, prior myocardial infarction, and endocarditis. Healthy volunteers and AD+CAA patients were matched based on age, sex, and ethnicity. Human cardiac specimens were used for biochemical and histological characterization.

### Statistical methods and analysis

Data are expressed as means±SEM. A two-tailed Student’s t-test determined the statistical significance between the two groups. When comparing multiple groups, data were analyzed using a one-way or two-away analysis of variance (ANOVA) test, followed by Tukey’s post hoc test. Outliers were identified and excluded from the original dataset using the ROUT (Q = 1%) method with GraphPad Prism 9.1.0 software. Differences between groups were considered statistically significant when p≤0.05. All data were analyzed and graphically represented using GraphPad Prism software version 9 (GraphPad, La Jolla, CA).

## 3. Results

### 3.1. AD pathology promotes cardiac dysfunction, Aβ accumulation, and misfolding of serum amyloid A3 in the heart of Tg2576 mice

To assess the impact of AD amyloid pathology on cardiac tissue physiology and systolic activity, we conducted echocardiographic analyses in equal numbers of male and female Tg2576 transgenic mice, and age-matched WT littermates, both before and after the development of cerebral Aβ pathology (at 4 and 8, or 13 months, respectively). Bi-dimensional M-mode tracings revealed a significant decrease in left ventricular ejection fraction (EF%) and fractional shortening (FS%) percentage in 13-month-old Tg2576 mice compared to WT animals (Fig. 1A-B, and other cardiac echocardiographic parameters in Supplementary Table 1). Radial and longitudinal strain/strain rates were also analyzed in both animal groups. Tg2576 exhibited global (average of 6 LV segments) and anterior-apical longitudinal strain impairment compared to WT mice, with severe longitudinal contractility dysfunction that affects the anterior-basal segment (Supplementary Fig. 2). Younger Tg2576 cohorts (at 4 or 8 months) did not exhibit significant changes in cardiac physiology (Supplementary Fig. 3), despite the presence of Aβ deposits in the brain starting at 8 months (Supplementary Fig. 5). Furthermore, 13-month-old transgenic mice displayed notable left ventricle enlargement, evidenced by increased left ventricular end-systolic diameter, leading to elevated left ventricular end-systolic volume (LVESV, Supplementary Table 1). The gravimetric analysis confirmed a significant increase in the heart weight/body weight ratio in 13-month-old Tg2576 mice compared to the WT group (Fig. 1C), further corroborated by the increased cardiomyocyte cross-sectional area assessed through WGA staining (Supplementary Fig. 1). Cardiac dysfunction in the Tg2576 mice coincided with a marked increase in myocardial interstitial fibrosis, as evidenced by Picrosirius red staining (Fig. 1D-E), accompanied by amyloid β accumulation in Tg2576 myocardial parenchyma (Fig. 1L) that contributes to a cardiac stiffness enhancement. Immunostaining results were confirmed via WB analysis, which shows high levels of fibronectin, a key component of the extracellular matrix (ECM) ^30^, in the Tg2576-AD mouse heart (Supplementary Fig. 1). However, in line with another recent AD study ^31^, no significant changes in cardiac MMP-2 levels between the WT and transgenic AD mice were observed (Supplementary Fig. 1).

**Figure 1.**
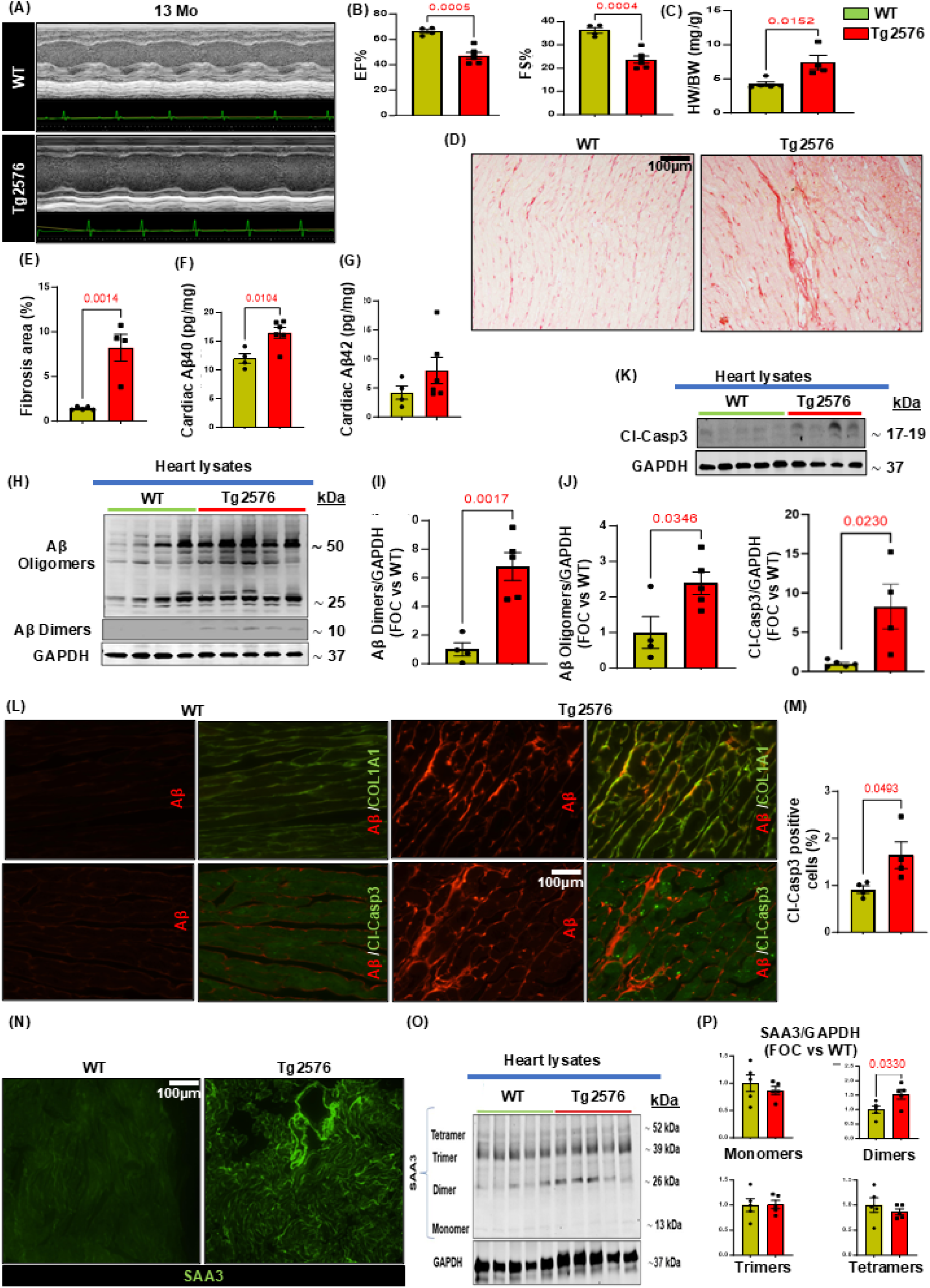
Cardiac physiological impairment, myocardial Aβ aggregates accumulation, cardiac cell apoptosis and Serum amyloid A3 (SAA3) deposition in the heart of Tg2576 AD mice. (**A**) M-mode representative images of echocardiographic analysis in 13-month-old WT and Tg2576 mice. (**B**) Left ventricle (LV) function is impaired in the AD model, as evidenced by a significant decline both in ejection fraction (EF%) and fraction shortening percentage (FS%) showed by the Tg2576 group compared to WT mice age-matched (13-months-old). (**C**) The heart weight/body weight ratio is significantly increased in Tg2576 mice compared to the WT group. (**D-E)** Representative images (left panels) and quantitative data (right panels) showing the percentage (%) of fibrotic area in cardiac sections from WT and Tg2576 mice, assessed via Picrosirius red staining (scale bar 50μm). (**F-G**): Aβ40 (F) Aβ42 (**G**) cardiac levels (pg/mg of proteins), assessed by ELISA assay, in total cardiac lysates from WT and Tg2576 mice. (**H-J**): Representative immunoblots (left panels) and densitometric quantitative analysis (right panels) showing protein levels of Aβ dimers, oligomers (**I-J**) in total cardiac lysates from WT and Tg2576 mice. GAPDH levels were used as a loading control. (K): Representative immunoblots (upper panels) and densitometric quantitative analysis (lower panels) showing protein levels of Cleaved-Caspase3 (Cl-Casp3) in total cardiac lysates from WT and Tg2576 mice. GAPDH levels were used as a loading control. (**L**): Representative digital images (upper panels, scale bar 100μm) showing collagen type I fibers (COL1A1, in green) and Aβ (in red) and representative digital images (lower panels; scale bar 100μm) and quantification (**M**) showing Cleaved-Caspase3 (Cl-Casp3, in green) and Aβ (in red) in cardiac sections from WT and Tg2576 mice. (**N**): Representative WT and Tg2576 cardiac sections stained for serum amyloid A3 (SAA3, in green, scale bar 100μm). (**O-P**): Representative immunoblots (**O**) and densitometric quantitative analysis (**P**) showing levels of different aggregates of SAA3 in total cardiac lysates from WT and Tg2576 mice. GAPDH levels were used as a loading control. n=4-6 mice/group. Data are presented as a mean±SEM. *P<0.05 and vs WT. Student t-tests were performed between the groups.

Using peptide-specific ELISA assays, we confirmed a significant increase in Aβ40 levels in 13-month-old Tg2576 cardiac tissue compared to WT mice, while no substantial changes in Aβ42 levels were noted between the two groups (Fig. 1F-G). These findings align with the age dependent Aβ pathology in the Tg2576 mouse model, characterized by higher abundance and progressive increase in cerebral Aβ40 levels compared to Aβ42^32^. Western blot analysis further validated the cardiac Aβ40 deposition detected by ELISA, confirming a significant increase in Aβ dimers and oligomers in total heart lysates of 13 months old Tg2576 mice compared to WT littermates (Fig. 1H-I-J). Immunofluorescence staining revealed amyloid aggregates accumulating in interstitial spaces between cardiomyocytes, alongside collagen fibers, corroborating the myocardial interstitial fibrosis observed via Picrosirius red staining (Fig. 1L, top panels, and Supplementary Fig. 6B). Besides Aβ aggregates, reports have demonstrated amyloid precursor protein (APP) expression in peripheral organs, including the myocardium, suggesting the potential production of Aβ outside the brain parenchyma^9,33,34^. Consistent with this, but also with the fact that the prion protein promoter may be expressed in other innervated tissue, including in the heart, we observed APP upregulation in the plasma and cardiac tissue of Tg2576 mice (Supplementary Fig. 8). Aβ challenge has been linked to increased apoptosis in various cell types, including cardiac cells, and in AD brains^35,36^. Indeed, analysis of cardiac tissue by western blotting and immunohistochemistry revealed elevated levels of cleaved-caspase 3 in Tg2576 cardiomyocytes (Fig. 1K, 1L bottom panels, 1M).

As AD pathology is characterized by progressive protein misfolding, seeding processes may be initiated by different amyloids interacting with each other^37^. Therefore, the accumulation of misfolded Aβ oligomers in the heart tissue may contribute to increased fibrosis and to the seeding of other amyloid peptides, such as serum amyloid A (SAA) proteins. Although high levels of SAAs typically do not lead to fibrillar deposits, chronic inflammatory states or seeding processes (e.g., promoted by Aβ) may induce the accumulation of SAA peptides in peripheral tissues^38,39^. Indeed, we observed Aβ aggregates around cardiac vessels in aged Tg2576 mice, accompanied by infiltration of the vascular basal lamina (Supplementary Fig. 4, Supplementary Fig. 6, *bottom panels*, and Supplementary Fig. 7D), alongside increased serum amyloid A3 (SAA3) vascular deposits (Fig. 1N). Immunoblot analysis confirmed increased SAA3 dimers in the cardiac tissue of Tg2576 mice (Fig. 1O-P). Overall, these findings indicate that AD pathology detrimentally affects systolic function and cardiac tissue structure. Progressive interstitial and perivascular Aβ deposits contribute to cardiac cell apoptosis and aggregation of serum amyloid proteins, culminating in cardiovascular dysfunction. Notably, 8-month-old Tg2576 mice exhibited evident amyloid aggregate accumulation in the prefrontal cortex vasculature and cerebral parenchyma (Supplementary Fig. 5), consistent with literature describing early Aβ deposition and CAA in the Tg2576 brain^32^. However, no significant changes were observed in Aβ accumulation in the hearts of 8-months old Tg2576 mice compared to the older transgenic cohort (Supplementary Fig. 5 and Supplementary Fig. 6), confirming that Aβ pathology starts in the brain of these mice, followed by a later increase in cardiac Aβ levels.

### 3.2. Neurotrophic signaling is impaired in the heart of AD transgenic mice

AD pathology and Aβ deposition are known to profoundly disrupt neurotrophins production in the brain, contributing to the progressive loss of key neuromodulators, including NGF and BDNF, ultimately leading to cognitive decline and dementia^40,41^. However, whether AD pathology can modulate peripheral neurotrophic activity and cardiac innervation remained to be elucidated. Therefore, we explored the regulation of neuro-signaling modulators and innervation factors in both cerebral cortex and cardiac lysates of 13-month-old Tg2576 mice. Immunoblotting analysis revealed a marked decrease in both cerebral and cardiac levels of NGF and BDNF expression in the Tg2576 group compared to WT littermates (Fig. 2A-B). This reduction in neurotrophins levels observed in transgenic mice was accompanied by a significant decrease in the adrenergic marker (DβH) and the neuronal fibers regeneration marker, GAP-43 (Fig. 2A-B). Intriguingly, immunohistochemistry analysis corroborated a notable loss in cardiac nerve fiber density in Tg2576 mice compared to WT animals, including reductions in sympathetic and regenerating nerve endings labeled with TH and GAP-43 markers, respectively (Fig. 2C-D-E and Supplementary Fig. 7A-C). These findings suggest that AD amyloid pathology may have detrimental effects on peripheral neurotrophic pathways, thereby impacting the peripheral nervous system and cardiac innervation.

**Figure 2.**
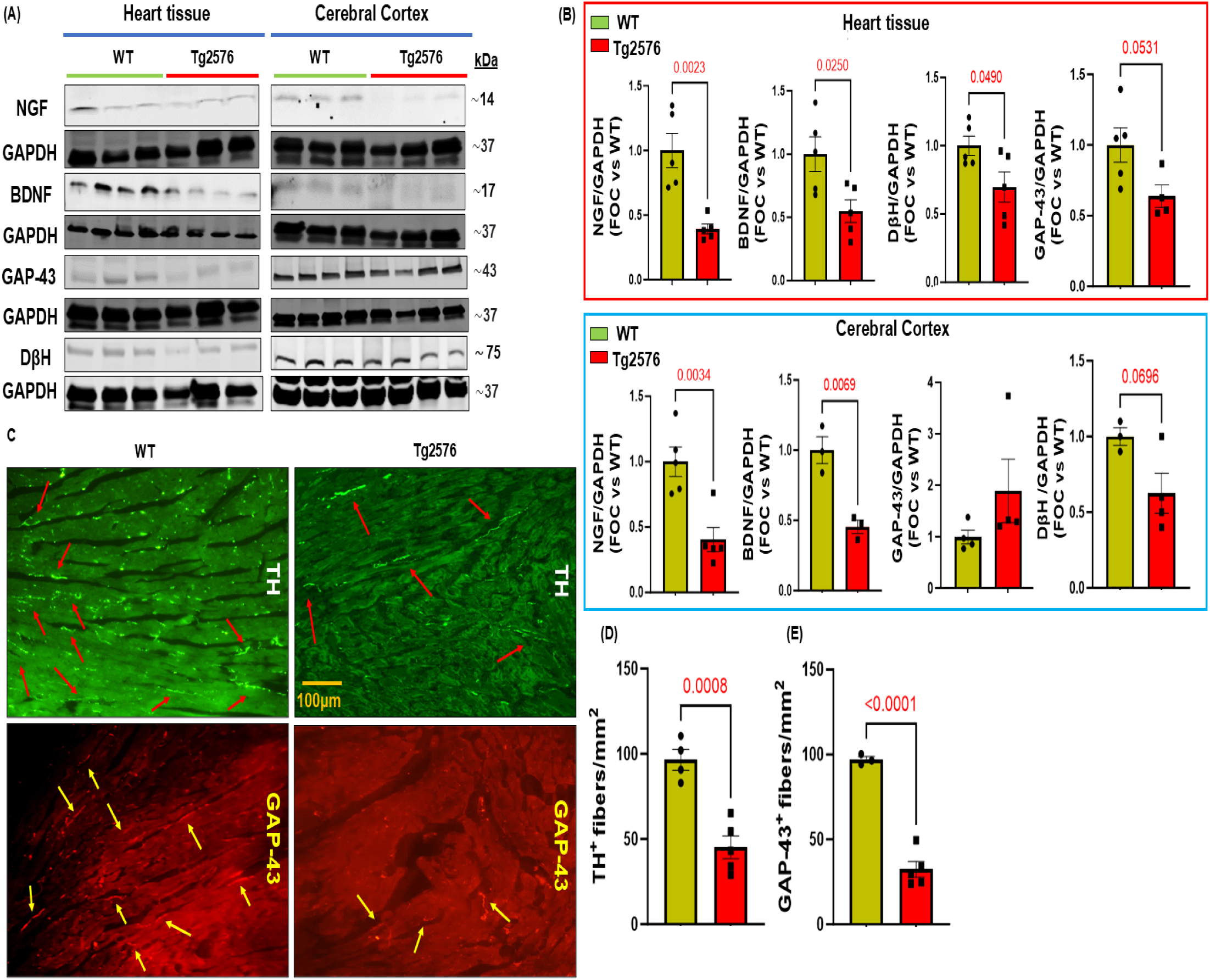
Alzheimer’s disease pathology negatively impacts the neuro-signaling pathway in the brain/heart axis of Tg2576 mice. (**A**-**B**): Representative immunoblots (**A**) and densitometric analysis (**B**) showing levels of NGF, BDNF, GAP-43, and dβh in total heart tissue and cerebral cortex lysates from WT and Tg2576 mice. GAPDH levels were used as a loading control. (**C**-**E**): Digital images (**C**, scale bar 100μm) and quantifications (**D**-**E**) showing cardiac adrenergic nerve fibers, labeled with anti-tyrosine-hydroxylase (TH, in green) (**D**), and cardiac regenerating nerve endings, labeled with anti-neuronal regeneration marker (GAP-43, in red) (**E**), in cardiac sections from WT and Tg2576 mice. n=3-5 mice/group. Data are presented as a mean±SEM. *P<0.05 and vs WT. Student t-tests have been performed between the groups.

### 3.3. Aβ oligomers impair neurotrophins production in hiPSC-CMs, AC16 cardiomyocytes and neuroblastoma cells

Our *in vivo* investigations revealed a notable decrease in neurotrophic markers and nerve fibers within the cardiac tissue of Tg2576 mice. To ascertain whether this loss of NTFs can be caused by the Aβ aggregates present in the heart, we evaluated the impact of Aβ40 oligomers, recognized as the most toxic amyloid aggregates and preferentially enriched in the AD brain vasculature, on the production of neurotrophins and neuronal regeneration markers in both human iPSC-CMs and AC16 cardiomyocytes (Fig. 3 and Fig. 4). Aβ40 oligomers significantly reduced BDNF expression in hiPSC-CMs and AC16 cardiomyocytes (Fig. 3E-F and Fig. 4). In addition, the effects of Aβ42 oligomers, the main toxic species coming in contact with neurons in the brain parenchyma, were examined on neuronal SHSY-5Y cells. Following a 16-hour incubation with Aβ oligomers, a significant reduction in protein levels of GAP-43 and BDNF was observed in neuronal cells, with no substantial changes in NGF expression (Fig. 3A-D). These findings support the hypothesis that Aβ toxic aggregates contribute to the impairment of the cardiac neuro-signaling and neural regeneration pathways by diminishing the production of BDNF in both neurons and cardiomyocytes and reducing levels of GAP-43 in neurons.

**Figure 3.**
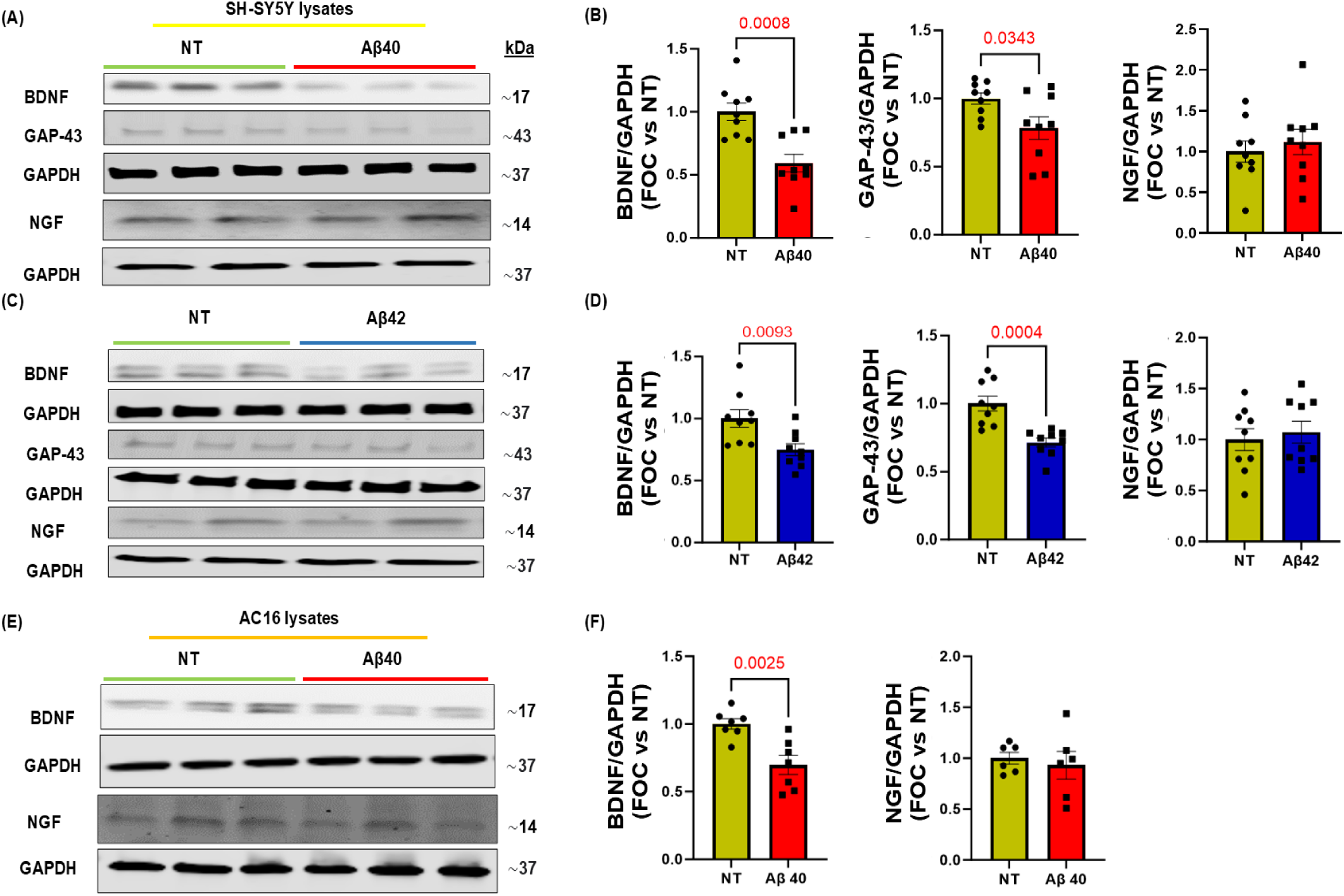
Effects of human Aβ40 or Aβ42 oligomers on neurotrophic factors in neuronal cells (SH-SY5Y) and human cardiomyocytes (AC16). (**A**-**D**): Representative immunoblots (**A**-**C**) and densitometric analysis (**B**-**D**) of multiple independent experiments to evaluate BDNF, NGF, and GAP-43 protein levels (**B**-**D**) in human neuroblastoma cells (SH-SY5Y) stimulated with human Aβ40 or Aβ42 oligomers for 16 h. GAPDH levels were used as a loading control. (**E**-**F**): Representative immunoblots (**E**) and densitometric quantitative analysis (**F**) showing protein levels of BDNF and NGF in total protein lysates from human cardiomyocytes (AC16) stimulated with human Aβ40 oligomers for 16 h. GAPDH levels were used as a loading control. n=6-9. Data are presented as a mean±SEM. *P<0.05 and vs NT. Student t-tests have been performed between the groups. (NT= Not Treated).

**Figure 4.**
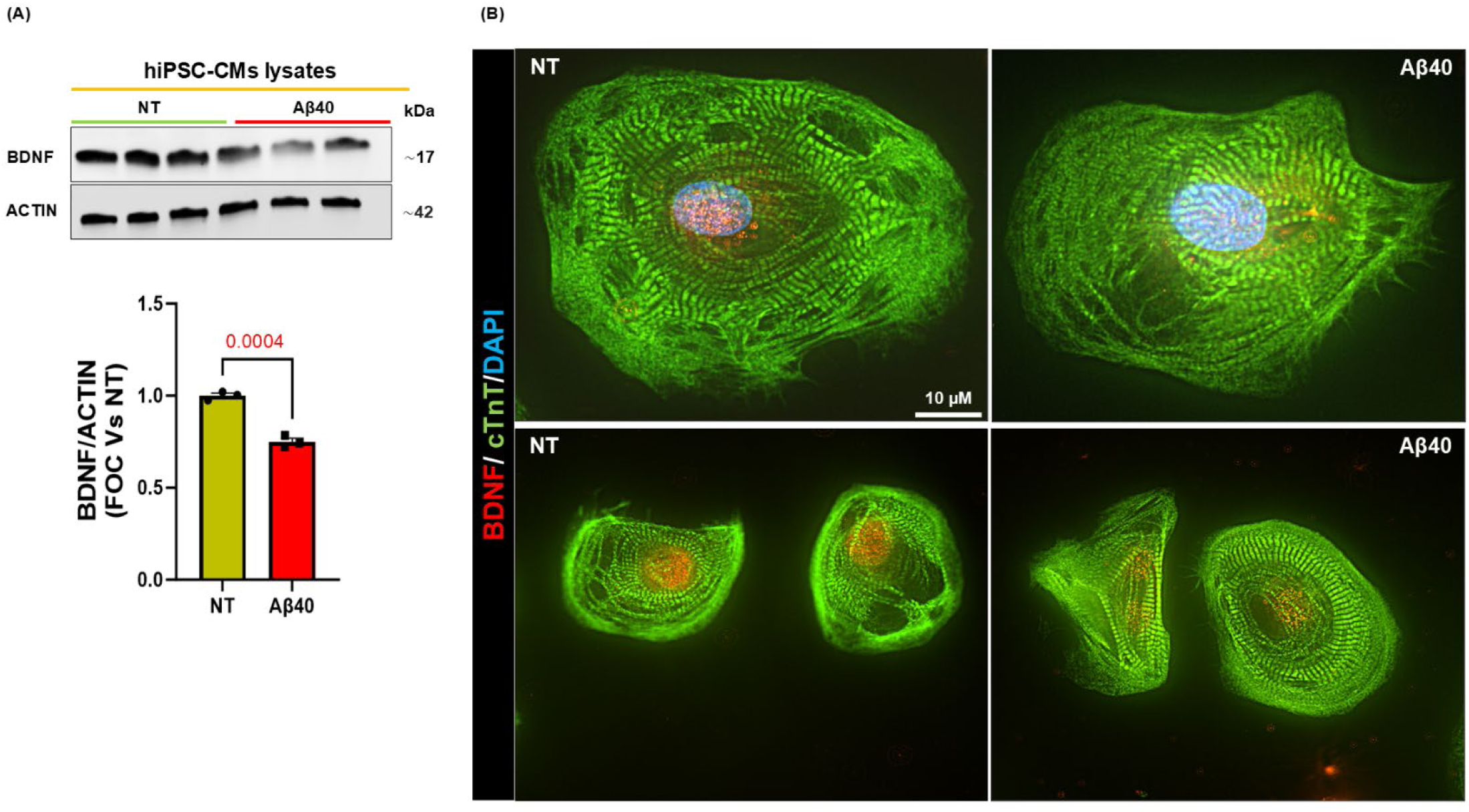
Aβ40 oligomers induce BDNF downregulation in human-induced pluripotent stem cell-derived cardiomyocytes (hiPSC-CMs). (**A**) Representative immunoblots (upper panels) and densitometric quantitative analysis (lower panel) showing protein levels of BDNF in total protein lysates from hiPSC-CMs stimulated with human Aβ40 oligomers for 20h. ACTIN levels were used as loading control. (**B**) Representative digital images (scale bar 10μm) of hiPSC-CMs stained with BDNF (red), cardiac Troponin T (cTnT, in green), and DAPI (blue). n=3. Data are presented as a mean±SEM. *P<0.05 and vs NT. Student t-tests have been performed between the groups. (NT= Not Treated).

### 3.4. Neurotrophic factors are reduced in the heart of Alzheimer’s disease patients with cerebral amyloid angiopathy

To confirm the clinical and translational relevance of our findings in human AD, we investigated the cardiac neurotrophic profile in LV biopsies obtained from patients affected by AD along with CAA. Our study cohort comprised individuals clinically diagnosed with AD, confirmed through anatomopathological examination, excluding cases affected by other conditions that could significantly impair cardiac function, such as high blood pressure, secondary amyloidosis, stroke, brain and cardiac neoplastic conditions, prior myocardial infarction, and endocarditis. Healthy volunteers and AD+CAA patients were matched based on age, sex, and ethnicity (Table 1). Our analysis revealed significantly reduced levels of BDNF, downregulation of the sympathetic marker tyrosine hydroxylase (TH), and a decline in GAP-43 protein expression in the heart of AD+CAA subjects, confirming the deterioration of neurotrophic factors and the neuro-signaling pathway in the human AD heart (Fig. 5A-B), with enormous clinical implications. Immunostaining analysis showed amyloid-β deposits in the myocardial parenchyma of human AD patients with CAA (Fig. 5C). In aggregate, these findings demonstrate a myocardial neuro-signaling impairment associated to Aβ deposition in human AD hearts, supporting our experimental *in vitro* and *in vivo* results.

**Table 1.** Anatomopathological and demographic characteristics of the selected clinical cohort. *P values of the patients age = 0.1253 vs CTRL. Student t-tests have been performed between the groups. **Abbreviations**: **CF**= Caucasian Female, **CM**= Caucasian Male, **Ctrl**= Control, **AD**= Alzheimer’s Disease, **CAA**= Cerebral Amyloid Angiopathy.

| Age | Ethnicity /Gender | Group | Associated Diseases and Comorbidities |
| --- | --- | --- | --- |
| 53 | CM | CTRL | Adenocarcinoma, Status post-Whipple procedure and liver transplantation, Hydroceles, Pulmonary vascular congestion, Hepatic congestion |
| 75 | CM | CTRL | Renal collecting duct carcinoma, Status post-Whipple procedure with gastric bypass and cholecystectomy, Mild cardiomegaly, Aorta Atherosclerosis, Testicular atrophy |
| 81 | CM | CTRL | Prostatic adenocarcinoma, Bilateral end-stage lung with honeycombing fibrosis (idiopathic pulmonary fibrosis with bronchiectasis by history), Cardiomegaly with left ventricular myocardial hypertrophy, Hepatomegaly, Hepatitis, Acute tubulointerstitial nephritis |
| 90 | CF | CTRL | Bilateral renal atrophy with mild nephrosclerosis and cortical cysts, Generalized atherosclerosis, Chronic esophagitis, Cardiomegaly with left ventricular myocardial hypertrophy, Hepatic centrilobular congestion, Chronic thyroiditis |
| 74 | CF | AD+CAA | Clear cell renal cell carcinoma, Cardiomegaly with left ventricular myocardial hypertrophy, Atherosclerosis, thoracic and abdominal aorta; Pulmonary vascular congestion; Hepatic atrophy; Status post thyroidectomy; Status post hysterectomy and bilateral salpingo-oophorectomy |
| 85 | CM | AD+CAA | History of prostatic carcinoma, status post prostatectomy, Aortic atherosclerosis, Left ventricular myocardial hypertrophy, Pulmonary fibrosis and bronchiolar metaplasia, Chronic esophagitis, Benign renal cysts, bilateral, Pancreatic and Testicular atrophy, Thyroid Nodular goiter, Status post cholecystectomy and Status post appendectomy |
| 86 | CF | AD+CAA | Acute bronchopneumonia, Mild cardiomegaly with left ventricular myocardial hypertrophy, Aortic atherosclerosis, Benign endometrial polyp, Nodular goiter, Osteopenia, Hepatic chronic passive congestion, Enchondroma, lumbar vertebra, Renal cortical cysts, History of plasmacytoma |
| 87 | CM | AD+CAA | Acute bronchopneumonia; Pulmonary interstitial fibrosis, Cardiomegaly with left ventricular myocardial hypertrophy, Aortic atherosclerosis, Chronic pyelonephritis, Bilateral renal atrophy, Hepatic atrophy, Cholelithiasis, Parathyroid adenoma, S/P radical prostatectomy, history of prostate cancer, S/P cardiac pacemaker placement |
| 87 | CM | AD+CAA | Acute bronchopneumonia, Aortic atherosclerosis, Hepatic chronic passive congestion, Prostatic glandular hyperplasia, Testicular atrophy, Mild chronic thyroiditis, Status post cardiac pacemaker, Diverticulosis, Benign renal cortical cyst, right |
| 89 | CF | AD+CAA | Acute bronchopneumonia, Calcified granuloma, Aortic atherosclerosis, Pleural fibrotic plaque, Mild calcification, aortic valve, Atherosclerotic stenosis, Cardiomegaly, Hepatic atrophy, Mild chronic lymphocytic thyroiditis, Status post hysterectomy and bilateral salpingo-oophorectomy |
| 90 | CF | AD+CAA | Pancreatic ductal adenocarcinoma, Cardiomegaly, Atherosclerosis, thoracic and abdominal aorta, Emphysema, Diverticulosis, Hashimoto's thyroiditis, Status post supracervical hysterectomy, Benign atrophic and fibrotic breast tissue, Osteopenia |

**Figure 5.**
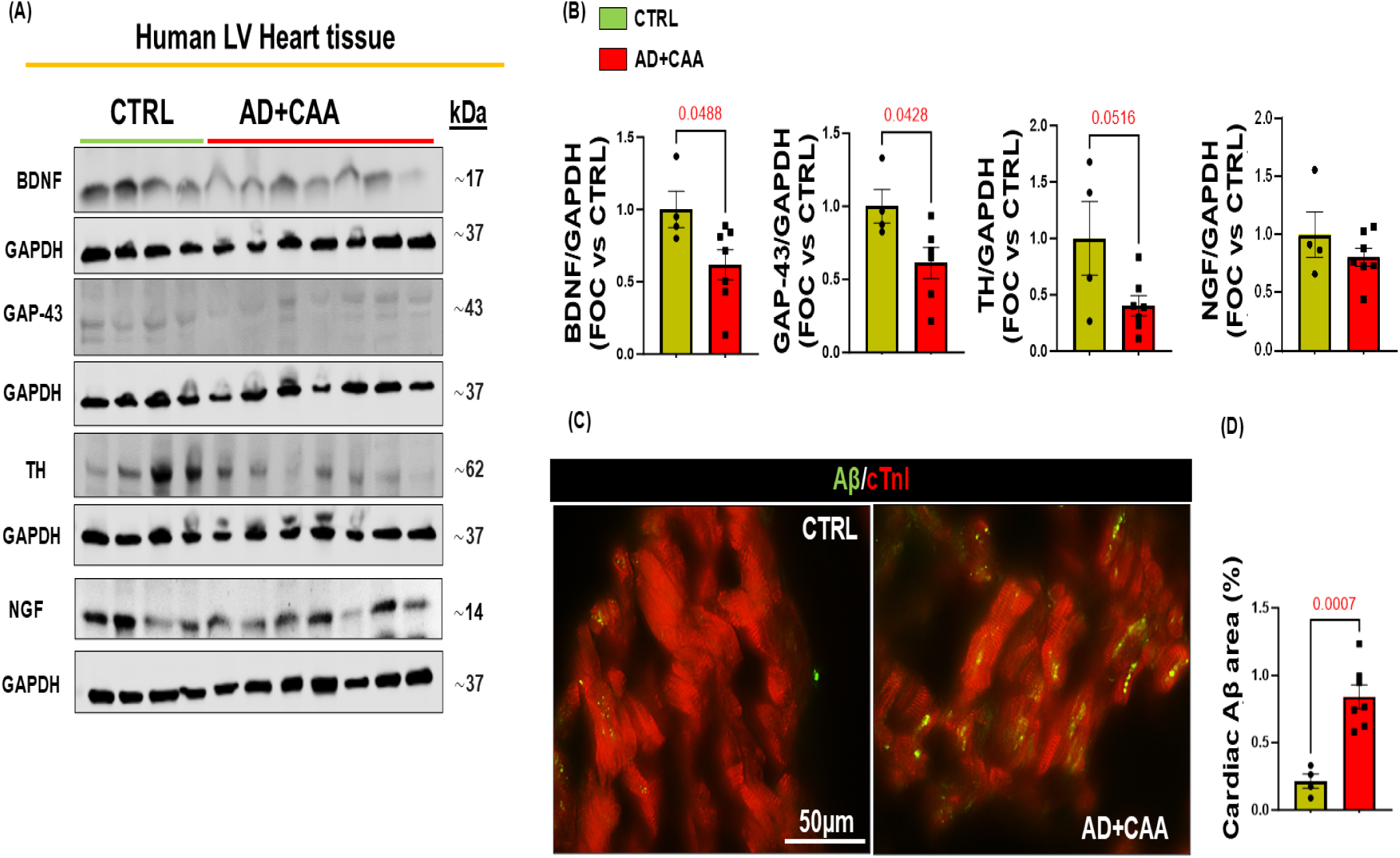
Myocardial neuro-signaling impairment and Aβ deposits in human post-mortem cardiac tissue of AD subjects. (**A**-**B**): Representative immunoblots (**A**) and densitometric analysis (**B**) showing reduced levels of BDNF, GAP-43, TH, but not NGF in total human LV lysates of patients with AD+CAA compared to healthy donors (CTRL). GAPDH levels were used as a loading control. (**C-D**): Representative digital images (**C**, scale bar 50μm), and quantification (**D**) of the LV sections from CTRL and AD+CAA subjects stained with Aβ aggregates (in green) and cardiac Troponin I (cTnI, in red). n=4-7 subjects/group. Data are presented as a mean±SEM. *P<0.05 and vs CTRL. Student t-tests have been performed between the groups.

### 3.5. Aβ40 oligomers impair BDNF/TrkB/CREB signaling in human cardiomyocytes

The downregulation in BDNF expression observed in our *in vitro* and *in vivo* experiments prompted us to investigate the mechanism by which Aβ affects the cardiac BDNF pathway. BDNF/TrkB/CREB signaling impairment is involved in age-related synaptic deterioration and neurodegenerative disorders, culminating in cognitive decline and neuronal loss. Indeed, AD patients exhibit significant depletion of BDNF expression in cerebrospinal fluid, post-mortem brain tissue, and bloodstream^42^. The detrimental impact of Aβ oligomers on BDNF exon IV transcript in neuronal cells is associated with decreased CREB levels, the critical mediator of BDNF transcription^43,44^. Consistent with these findings, here we observed a significant reduction in nuclear CREB levels in human cardiomyocytes following treatment with Aβ40 oligomers (Fig. 6A-C), suggesting that a reduction in CREB transcriptional function mediates both the reduced levels of BDNF and the impact on the other neurotrophic markers. These results confirm the critical role of the BDNF/CREB axis in AD cardiac pathology, unveiling an Aβ-dependent signaling mechanism which can explain the BDNF reduction we observed in AD. To confirm the protective effect of TrkB activation in our model, we stimulated human cardiomyocytes with the selective TrkB receptor agonist, LM22A-4^45^ (Fig. 6D-E), in the presence or absence of Aβ40 oligomers^46^. Remarkably, LM22A-4 stimulation reduced Aβ40-mediated caspase 3 activation, thereby mitigating apoptosis, and confirming the protective effects of BDNF signaling on cardiac cells (Fig. 6G-H). Conversely, ANA-12, a specific BDNF/TrkB receptor antagonist^47^ (Fig. 6D-F), exacerbated cardiomyocyte caspase 3 activation, (Fig. 6G-I), Overall, these findings elucidate the mechanisms and impact of the neurotrophic signaling deregulation induced by Aβ pathology in the heart (Supplementary Fig. 9), highlighting the protective role of BDNF/TrkB/CREB signaling in cardiomyocyte homeostasis.

**Figure 6.**
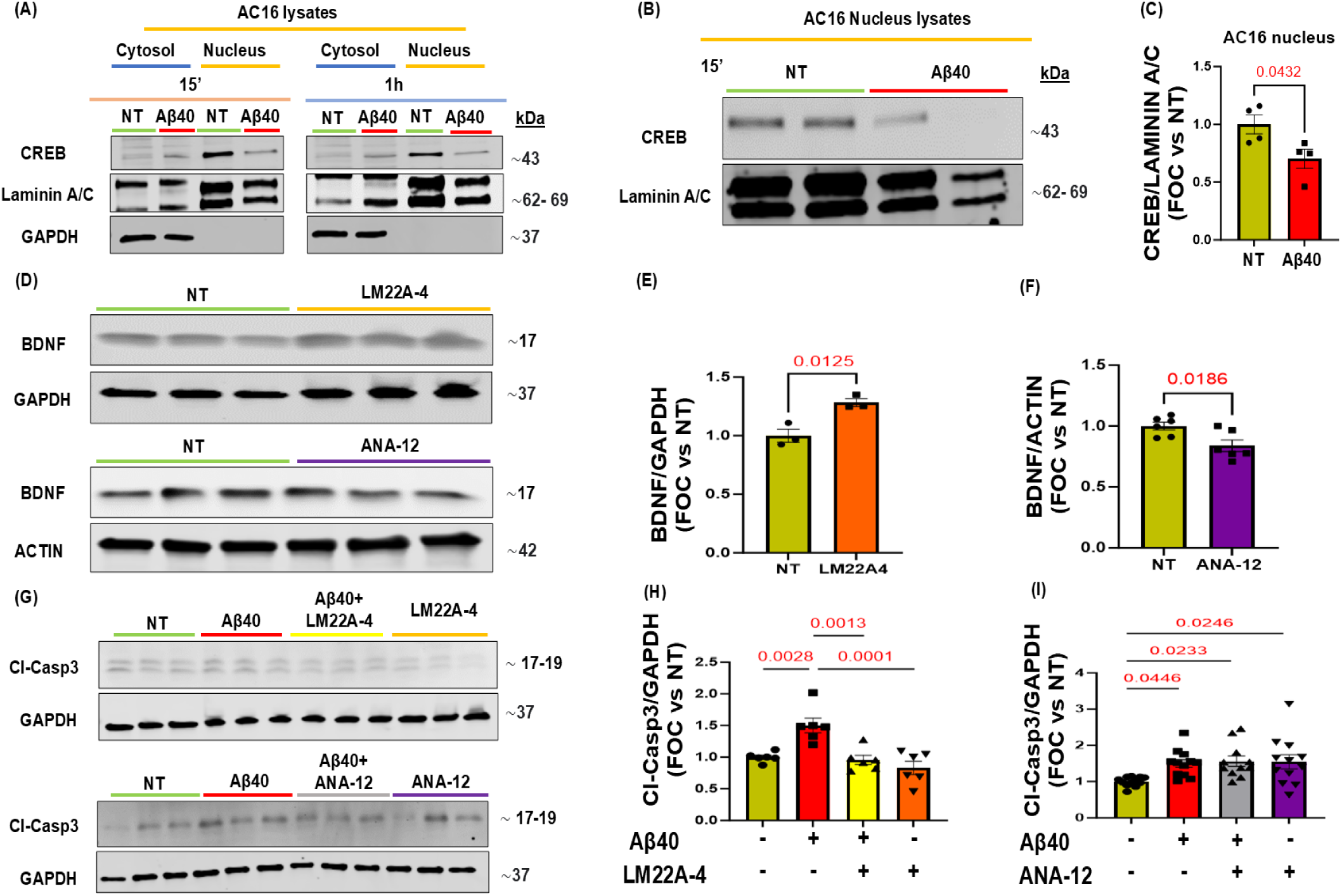
CREB signaling is impaired by Aβ40 oligomers and a TrkB agonist reduces Aβ40-mediated caspase 3 activation in human cardiomyocytes (AC16 cells). (**A**-**C**): Representative immunoblots (**A**-**B**) and densitometric quantitative analysis (**C**) of multiple independent experiments to evaluate CREB levels in nuclear extracts isolated from human cardiomyocytes (AC16) stimulated with human Aβ-40 oligomers for 15 minutes or 1h. LAMININ A/C levels were used as a nuclear loading control. (**D**-**F**): Representative immunoblots (**D**) and densitometric quantitative analysis (**E**-**F**) showing protein levels of BDNF in total protein lysates from human cardiomyocytes (AC16) stimulated with TrkB agonist LM22A-4 (100 nM) or TrkB antagonist ANA-12 (30 µM) for 16 h. GAPDH and ACTIN levels were used as loading controls. n=3-6. Data are presented as a mean±SEM. *P<0.05 and vs NT. Student t-tests have been performed between the groups. (**G**-**I**): Representative immunoblots (**G**) and densitometric quantitative analysis (**H**-**I**) of multiple independent experiments to evaluate Cleaved-Caspase 3 (Cl-Casp3) in AC16 cells stimulated with human Aβ-40 oligomers for 16 h. Before Aβ-40 oligomers treatment, cells were pretreated with either LM22A-4 (100 nM, top panel), or ANA-12 (30 µM, bottom panel) for 1 h. GAPDH levels were used as a loading control. n=6-12. One-way ANOVA with Tukey’s multiple comparisons post hoc test: *P<0.05 and vs NT. (NT= Not Treated).

## 4. Discussion

This study demonstrates that, similar to what was previously observed in the brain, Aβ deposition in the AD heart leads to neurotrophic signaling impairment, which contributes to increased cardiomyocyte apoptosis and severe cardiac morphological and functional impairment observed in the hearts of Tg2576 mice. Confirming the clinical validity of these findings, the neurotrophic changes and the presence of Aβ pathology were validated in human post-mortem heart samples from patients with AD+CAA. Our data suggest that cardiac amyloidosis (evidenced by the presence of Aβ and SAA3 oligomers) contributes to interstitial fibrosis and to the loss of neurotrophins, resulting in cardiac innervation deficits and neuro-signaling pathway impairment, which appear to be essential mediators of the observed cardiac physiological dysfunction. The heart-brain axis constitutes a complex and highly regulated bi-directional interconnection between the central and peripheral nervous systems.^48^. Neurodegenerative pathologies, such as AD, may have severe and debilitating effects on various peripheral organs, including the heart, through the modulation of both the central and peripheral nervous systems. However, the impact of AD pathology and Aβ accumulation on cardiac neuro-signaling pathways in AD patients and animal models has not been previously elucidated. An increasing number of studies are beginning to highlight the involvement of cardiac tissue in neurodegenerative diseases, revealing a compelling interplay between AD and cardiac disorders^49^, such as atrial fibrillation^50^, heart failure^51^, and coronary artery disease^34^. It has recently been shown that AD patients exhibit myocardial dysfunction following the deposition of Aβ aggregates in cardiac tissue ^9^, and the presence of cardiac N-terminal cleavage products of APP was correlated with levels of CAA and myocardial fibrosis^52^. In line with this, Sanna and coworkers described the presence of structural and functional abnormalities in the cardiac tissue of a small cohort of AD subjects^4^.

AD pathology has been shown to severely impact the production and release of the main NTFs, NGF and BDNF, in the brain^53,54^. These two neurotrophins are mainly involved in the maturation and differentiation of neurons, regulating neurogenesis and synaptogenesis^55,56^. The progressive decline in NTFs expression due to aging, exacerbated by AD pathology, may lead to an adverse remodeling of the whole neuro-signaling pathway, driving, together with other AD pathological factors such as amyloid oligomers, an impairment of the peripheral innervation^41^.

This study demonstrates the effect of Aβ pathology on cardiac physiology and heart dysfunction using a transgenic mouse model of AD with CAA, human cardiomyocytes in culture, validated also in hiPSC-CMs, and AD+CAA human post-mortem heart tissue. It describes the mechanisms underlying the impairment of peripheral innervation and the loss of neurotrophins in the AD heart and shows a causal role of Aβ oligomers in the impairment of neurotrophic signaling in cardiomyocytes. We report that Tg2576 mice show an age-dependent deterioration in cardiac function and a gradual replacement of viable myocardial with fibrotic tissue, resulting in interstitial fibrosis. This cardiac maladaptive remodeling, which develops after the brain Aβ deposition, is accompanied by an increased cross-sectional area alongside a significant accumulation of Aβ, particularly Aβ40, localized in cardiac perivascular and interstitial spaces with collagen deposits. Caspase 3 activation within cardiomyocytes is also increased, suggesting that Aβ aggregates in the heart tissue contribute to the development of interstitial fibrosis and apoptotic cardiac cell death. The preferential accumulation of Aβ40, rather than Aβ42, in Tg2576 mice hearts is in line with previous studies describing the progressive increase mostly of Aβ40 levels in both serum and brain parenchyma of Tg2576 mice during aging^32^. Additionally, the Aβ40 peptide is preferentially associated with vascular deposits and CAA in the human brain, resulting in degenerative phenotypes in cerebral endothelial cells ^36^, which culminate in vessel wall dysfunction^35^, contributing to BBB permeability and microhemorrhages^57^. This increased BBB permeability in AD brains may enhance the systemic diffusion of Aβ40 (including aggregated forms such as oligomers) through the blood and its accumulation in peripheral organs, including the heart. Indeed, Aβ40 is the most abundant peptide in the cerebrospinal fluid and blood of AD patients, and multiple biomarker studies have shown that this peptide does not decrease in peripheral organs and tissues with AD progression^58^. Conversely, as Aβ42 fibrils accumulate in the brain parenchyma, Aβ42 levels decrease in the cerebrospinal fluid and in the peripheral blood^59^.

It is known that Aβ oligomers may act as “seeds”^60^ to induce aggregation of other amyloids. Some of these aggregation species fuel cell stress and inflammation, resulting in degenerative processes in multiple cell types, including endothelial and cardiac cells, as our group and others have shown^61,62^. Here, we show that SAA3 also accumulates in the heart of Tg2576 mice, likely contributing to amorphous and misfolded protein accumulation and to progressive cardiac impairment. SAA3 is involved in vascular injury as a promoter of atherosclerosis, and it is a prognostic factor for coronary artery syndromes, also proposed as a novel inflammatory marker for ischemic heart disorders^63–65^. Notably, we observed SAA3 aggregates accumulate primarily at the cardiac vessel walls, together with Aβ deposits, suggesting that misfolded Aβ peptides or oligomers may diffuse through the vascular system, reaching the heart, with consequential seeding of other systemic amyloids around the myocardial vessels. Interestingly, no changes in cardiac physiology and no evidence of amyloid deposits were found in younger (4 or 8 months) Tg2576 mice. However, Aβ deposits were already detected around brain vessels in the prefrontal cortex of 8-month-old Tg2576 mice. Thus, it is reasonable to think that, in this model and presumably in AD patients, cardiac dysfunction may be subsequent to the progressive accumulation of amyloid aggregates in the brain, particularly as CAA, in and around the brain vessels. Given the extensive literature available on the vascular contributions to cognitive impairment (VCID) field^5,66^, we postulate that the cardiac disease and maladaptive heart remodeling resulting from progressive amyloid deposition in the heart also contribute to cognitive decline, thereby exacerbating AD pathology in a destructive feed-forward loop.

Notably, we observed pro-apoptotic markers (active caspase 3) in cardiomyocytes exposed to Aβ oligomers in vitro and in vivo, suggesting a direct effect of Aβ pathological aggregates on the impairment of cardiac cell function. Interestingly, caspase 3 activation is also associated with tau cleavage at position 421, which promotes increased tau phosphorylation^67^, suggesting possible downstream effects that may precipitate tau pathology, which was recently described in the heart by the Del Monte group^8,68^. Notably, we revealed a depletion in both adrenergic nerve endings and regenerated nerve terminal fibers in the transgenic AD mouse hearts. This myocardial denervation was associated with a reduction of NGF and BDNF at both the cerebral and cardiac levels, accompanied by significant impairment in the neuronal sprouting marker GAP-43. Importantly, post-mortem AD human hearts also exhibited a reduction in BDNF expression, as well as in the cardiac nerve fiber markers TH and GAP-43. In AD patients, the decline of neuromodulators in the brain is known to accelerate age-related synaptic loss, resulting in cognitive impairment^69,70^. NTFs are powerful biomolecules with significant pleiotropic effects involved in cellular growth and survival pathways in both the brain and peripheral cells/tissues, including endothelial cells, muscle cells, and the cardiovascular system^22^. Our finding that after treatment with Aβ oligomers, BDNF and GAP-43 expression in human neurons, endothelial cells, and both hiPSC-CMs-derived and immortalized cardiomyocytes were significantly reduced suggests a direct effect of Aβ in lowering NTFs not only in neuronal cells, and possibly peripheral neurons, but also in endothelial cells and cardiomyocytes. This reduction may cause AD patients to be more susceptible to developing cardiovascular disorders of various etiologies^71,72^.

To validate the mechanism inducing this cardiac Aβ-mediated neuro-signaling deterioration, we manipulated the BDNF signaling pathway pharmacologically in human cardiomyocytes. Cells treated with LM22A-4, a competitive agonist of the BDNF receptor TrkB, showed a reduction in Aβ40 oligomers-mediated caspase 3 activation, confirming a TrkB-dependent mechanism linking reduced BDNF to the Aβ-mediated cardiomyocyte toxicity. LM22A-4 was also reported to alleviate cardiac ischemia-induced apoptosis and to improve neuronal outgrowth and myocyte physiology with endothelial cell proliferation, thus arresting chronic heart failure^14^. Moreover, LM22A-4 showed effects similar to BDNF in preventing Aβ-induced neuronal cell death^45^. These beneficial effects suggest the initiation of a myocardial autocrine/paracrine protective pathway through TrkB activation, which may include the increased transcription of BDNF itself, via CREB activation^14^.

In line with this concept, we demonstrated a significant downregulation of CREB nuclear expression (necessary for its transcriptional function) in cardiomyocytes challenged with Aβ40 oligomers. CREB is the leading transcription factor activated upon BDNF-TrkB binding, which also promotes, in a feed-forward loop, the transcription of BDNF itself. Our results provide a mechanistic explanation of how Aβ oligomers can promote a CREB-mediated downregulation of BDNF expression in cardiomyocytes, uncovering a previously unknown molecular pathway underlying the AD-mediated cardiac neuro-signaling dysregulation. Notably, we demonstrated that the cardiac tissue of AD patients with CAA exhibits significantly reduced neurotrophic markers, including BDNF, together with Aβ deposits within the myocardial tissue. The presence of Aβ aggregates was also previously revealed in the human AD heart^9^, reinforcing the relevance of our findings.

To our knowledge, this is the first study that investigated the impact of Aβ pathology on the neurotrophic signaling pathway in the AD heart and its effect on cardiovascular physiology in an AD animal model, with a validation of the loss of neurotrophic signaling mediators in human AD post-mortem cardiac tissue. These findings uncover a previously unrecognized dysfunction in the BDNF/TrkB/CREB signaling pathway, driven by Aβ accumulation within the cardiac nervous system. This disruption highlights neurotrophic signaling as a promising and novel pharmacological target for combating AD-induced cardiac dysfunction. While further clinical and translational studies are needed to fully validate these conclusions, our results strongly suggest that treating AD-related heart failure (HF) may require a therapeutic strategy specifically aimed at the cardiac neuronal system, that goes beyond conventional HF pharmacotherapy.

This emerging perspective calls for deeper investigation into the neuro-cardiac axis in AD. Importantly, our study opens a new translational avenue that may help personalize AD therapy in subjects with cardiac pathology, as well as characterize AD as a multi-organ neurodegenerative syndrome.

### 4.1 Limitations

While our study provides valuable insights into the cardiac implications of AD, several limitations should be acknowledged. First, we did not assess left ventricular diastolic function in our in vivo model, which has been implicated in previous studies of AD patients demonstrating diastolic impairment correlated with cardiac electrical abnormalities. Of note, given the advanced age of the animals used in the study, some technical difficulties do not always allow for consistent measurement of E/A ratios in all animals, given that the E and A waves overlap at a higher heart rate, as already demonstrated^18^. Therefore, the E/E′ ratio is not always considered a trustworthy measurement to assess the diastolic function in aging in vivo models, as also reported in murine heart failure with preserved ejection fraction and in volume overload models^19^. Accordingly, diastolic dysfunction was computed by using the reverse peak option to evaluate reverse global longitudinal strain (GLS) and global radial strain (GRS) rates, which were derived from long-axis B-mode traces. Moreover, our focus on systolic function deterioration aligns with findings linking systolic impairment to cerebral hypoperfusion and emboli formation in AD patients, highlighting the connection between Aβ pathology and cardiovascular disease^73^. Additionally, due to the frailty of our AD mouse model and the desire to minimize stress-induced effects, we refrained from performing hemodynamic analysis, which could have provided further insights but may have impacted our animal population adversely^74^. Other neurotrophic factors, such as NT3 and NT4, may also contribute to cardiac innervation and function. Future investigations should explore the potential involvement of these factors in the AD myocardium to provide a comprehensive understanding of the cardiac neuro-signaling dysregulation. The potential of the BDNF/TrkB/CREB signaling reactivation to reverse or prevent cardiac dysfunction remains to be tested in vivo. Studies employing pharmacological strategies to increase cerebral amyloid clearance are also under development^75^, to determine if reducing brain amyloid pathology may ameliorate heart function. Additionally, generating AD mouse models with BDNF cardiomyocyte-specific deletion could significantly enhance future research approaches. However, the combined mutations may lead to early morbidity and reduced lifespan^76,77^, therefore careful experimental design and substantial investment in resources and technology will be needed. Furthermore, Aβ plaques exacerbate cellular stress and inflammation, driving degenerative processes across multiple cell types, including neurons and brain endothelial cells. This cascade worsens BBB breakdown and accelerates AD-induced maladaptive remodeling. In our study, we demonstrated that SAA3, a known inflammatory marker for ischemic heart disease^63–65^, predominantly accumulates in the cardiac vessel walls of Tg2576-AD mice, together with Aβ deposits. This suggests that misfolded Aβ peptides or oligomers may infiltrate the vascular system, reaching the heart and fostering the seeding of other systemic amyloid deposition around myocardial vessels. These findings highlight the need for further investigation into the inflammatory dynamics of Alzheimer’s heart pathology, an intriguing avenue for future research beyond the scope of this study. Lastly, the sample size of our human cardiac post-mortem specimens is limited, as obtaining human cardiac samples from AD patients without significant cardiovascular diseases is extremely challenging. Future, more comprehensive clinical studies are recommended to confirm AD cardiac dysfunction and validate the Aβ-mediated myocardial denervation observed in our study. Despite these limitations, our findings offer novel key insights into the pathophysiology of AD-related cardiac dysfunction and underscore the need for further research to improve the clinical management of AD patients.

## 5. Conclusions

In summary, our study elucidates a profound impact of AD amyloid pathology on cardiac function, highlighting the maladaptive cardiac remodeling characterized by amyloid deposition, fibrosis, and a previously unknown dysregulation of the neurotrophic signaling pathway. The study also uncovers a mechanism whereby Aβ-induced reduction in cardiac CREB activation impairs BDNF transcription, ultimately leading to myocardial denervation, pro-apoptotic signaling, and cardiac physiological dysfunction. These findings significantly contribute to our understanding of AD pathophysiology, particularly Aβ-mediated cardiac dysfunction. Moreover, our study identifies potential therapeutic targets within the neurotrophic signaling pathway to mitigate the detrimental effects of AD and amyloid pathology on the cardiovascular system, bridging the gap between molecular insights and potential for clinical translation. Finally, this study underscores the importance of considering cardiac complications in managing AD and offers promising avenues for future research and therapeutic development.

## 6. Acknowledgement

We acknowledge that Supplementary Figure 9 was created using BioRender.com. We thank the Brain and Body Donation Program Banner Sun Health Research Institute clinical biobank for providing human LV post-mortem tissue. We thank Life Science Editors for editing services.

## 8. Authors Contribution

AE and SF designed the study. AE performed and conceptualized the experiments and analyzed the data. AE wrote the paper draft and did the literature search. RPR, AC, and RVT performed experiments. SF provided relevant insights, scientific supervision, critically revised the manuscript and acquired funding. SJ revised the study, edited the manuscript, and offered additional literature searches.

## 9. Data availability

The corresponding author will share the data underlying this article upon reasonable request.

## Figures

**Supplemental Table 1.**
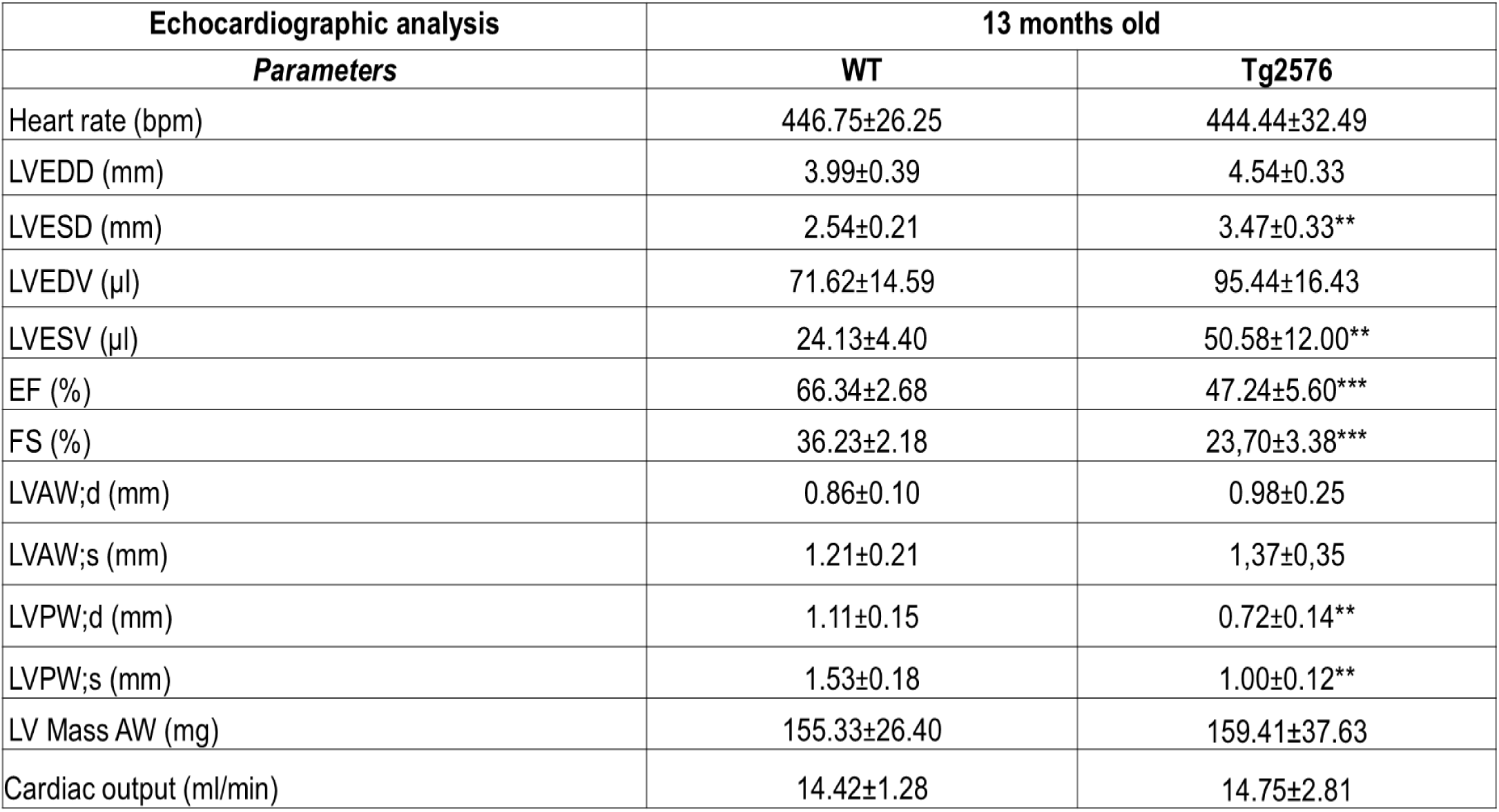
Echocardiographic evaluation. Conventional echocardiographic parameters were measured in 13-month-old WT and Tg2576 mice: heart rate, left ventricular (LV) end diastolic diameter (LVEDD), end systolic diameter (LVESD), end-diastolic volume (LVEDV), end systolic volume (LVESV), ejection fraction (EF), fractional shortening (FS), anterior wall in diastole (LVAW;d) and systole (LVAW;s), posterior wall in diastole (LVPW;d) and systole (LVPW;s), mass of the anterior wall (LV Mass AW), and cardiac output. n=4-6 mice/group. Data are presented as a mean±SEM. *P<0.05, **p<0.01, and ***p<0.001 vs WT. Student t-tests were performed between the groups.

**Supplemental Figure 1.**
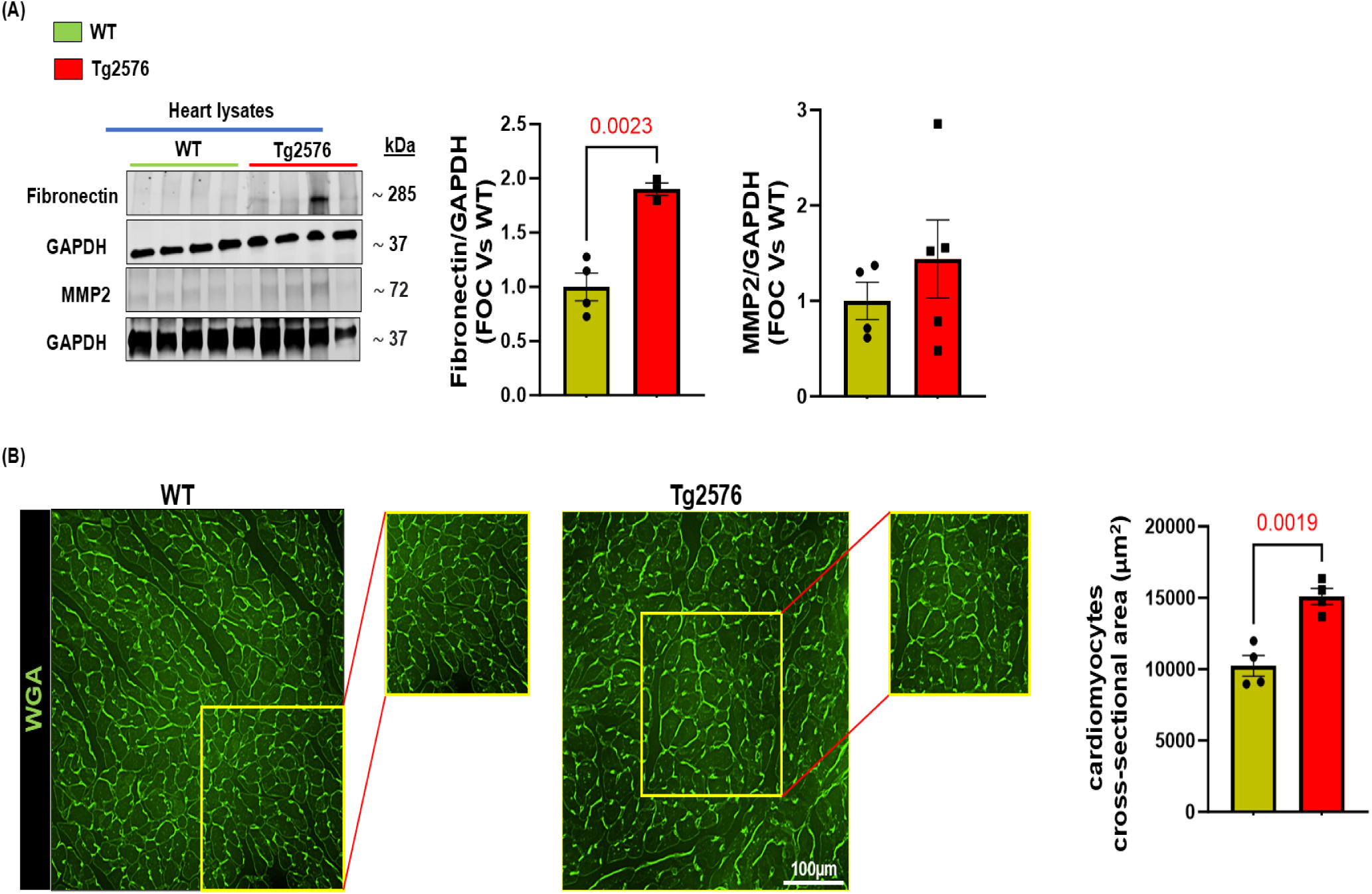
Cardiac adverse remodeling in 13-month-old Tg2576-AD mouse heart. (**A**) Representative immunoblots (left panels) and densitometric quantitative analysis (right panels) showing protein levels of fibronectin and MMP2 in total cardiac lysates from WT and Tg2576 mice. GAPDH levels were used as a loading control. (**B**) Representative digital images (left panels, scale bar 100μm), and quantification (right panel) of the cardiomyocyte cross-sectional area (CSA) in cardiac sections from WT and Tg2576 mice stained with WGA. n=4-5 mice/group. Data are presented as a mean±SEM. *P<0.05 and vs WT. Student t-tests have been performed between the groups.

**Supplemental Figure 2.**
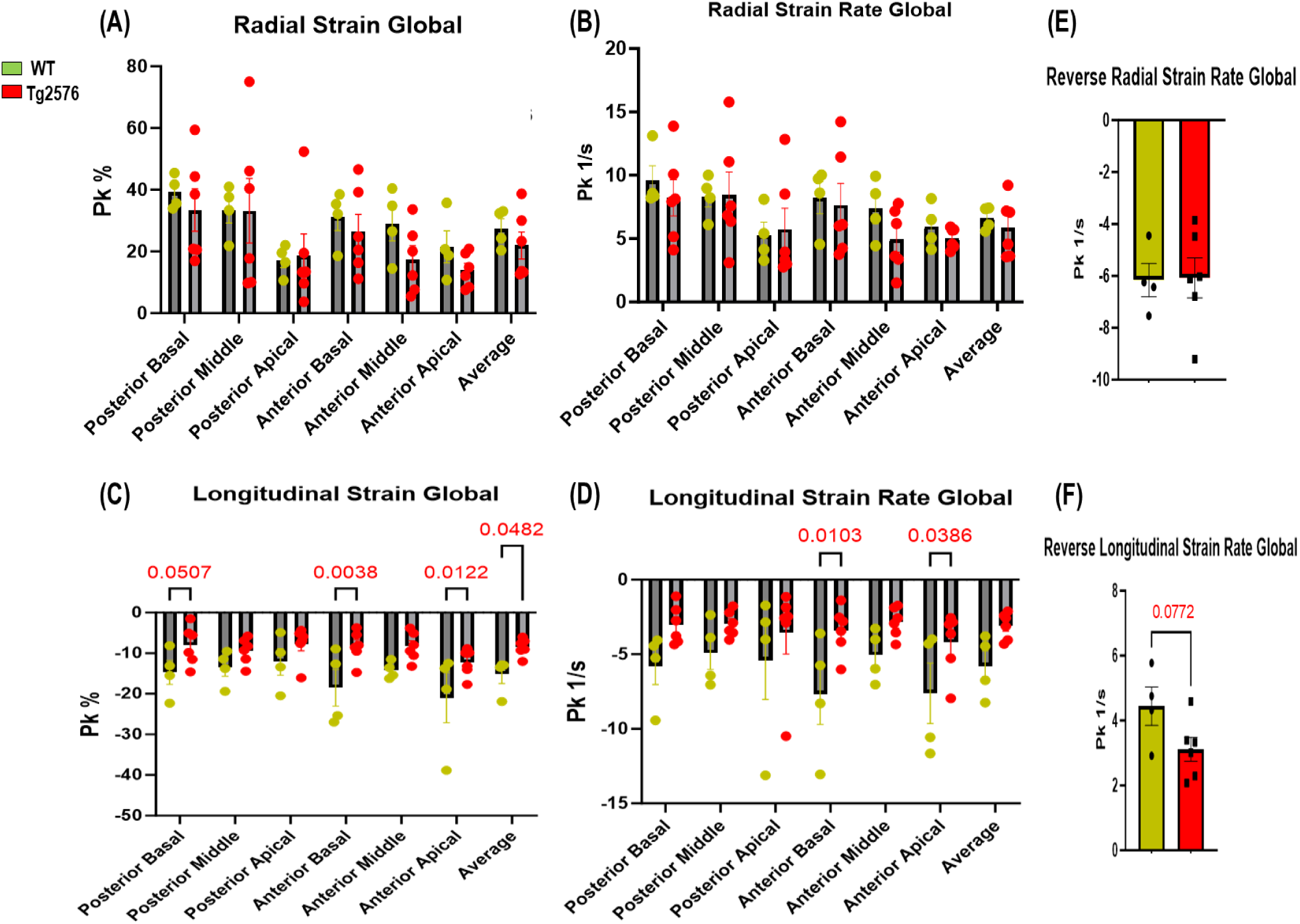
Longitudinal strain and strain rate LV contractility are impaired in Tg2576 AD mice. Radial and longitudinal strain or strain rate measured at six LV segments as well as their average(**A**-**D**), and reverse (diastolic) radial (**E**) and longitudinal (**F**) SR global in 13 months old WT and Tg2576 mice were evaluated using speckle-tracking-based strain echocardiography. Data are presented as a mean±SEM. *P<0.05 and vs WT. n=4-6 mice/group. Two-way ANOVA with Tukey’s multiple comparisons post hoc test (**A**-**D**), and student t-tests (**E**-**F**) have been performed between the groups.

**Supplemental Figure 3.**
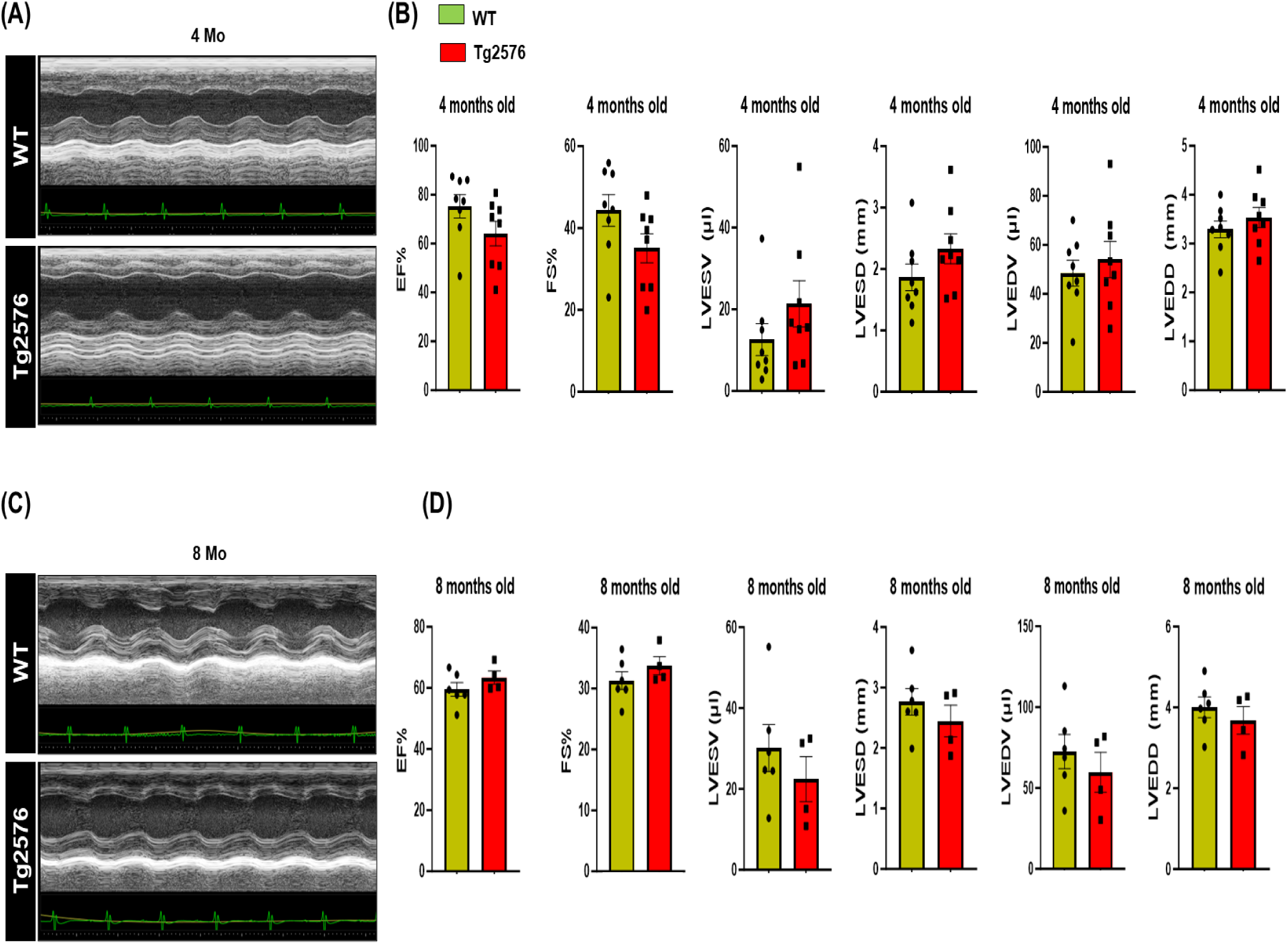
Cardiac function is not affected in 4 or 8 months old Tg2576 mice. (**A**-**C**) Representative M-mode echocardiography images of WT and Tg2576 mice show no difference between the experimental groups. (**B**-**D**) Ejection fraction (EF%) and fractional shortening (FS%) percentage, left ventricular end-systolic and diastolic diameters (LVESD, LVEDD), and left ventricular end-systolic and diastolic volumes (LVESV, LVEDV) of age-matched (4 and 8-month-old) Tg2576 mice and WT littermates. n=4-8 mice/group. Data are presented as a mean±SEM. *P<0.05 and vs WT. Student t-tests have been performed between the groups.

**Supplemental Figure 4.**
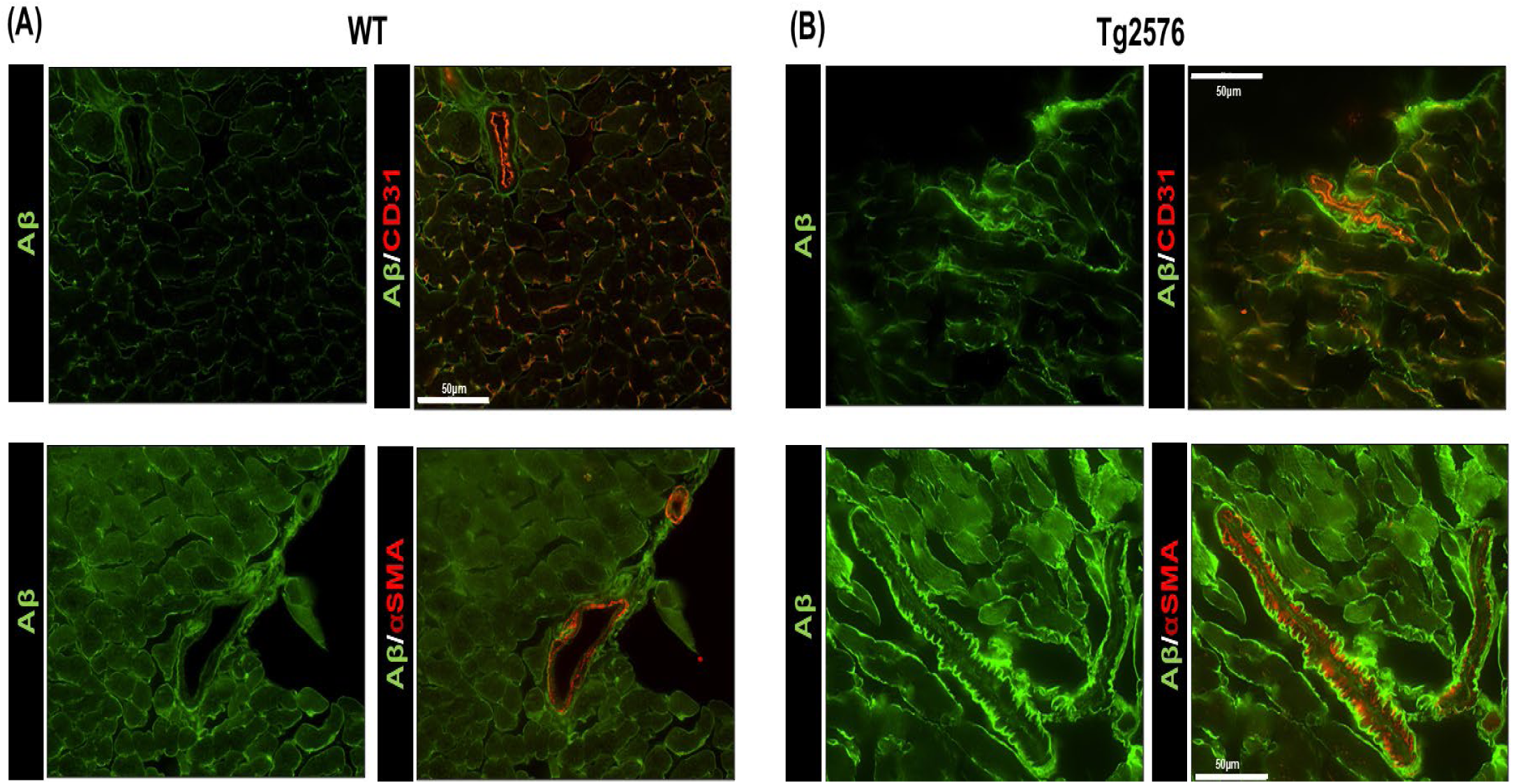
Amyloid-β surrounds cardiac vessels and infiltrates the vascular basal lamina. (**A**-**B**) Representative digital images (upper panels, scale bar 50μm) showing vessels (endothelial cells stained with CD31, in red) and Aβ (in green), and representative digital images (lower panels; scale bar 50μm) showing α-smooth muscle actin (αSMA, staining vascular smooth muscle cells, in red) and Aβ (in green) in cardiac sections from 13 months old WT and Tg2576 mice.

**Supplemental Figure 5.**
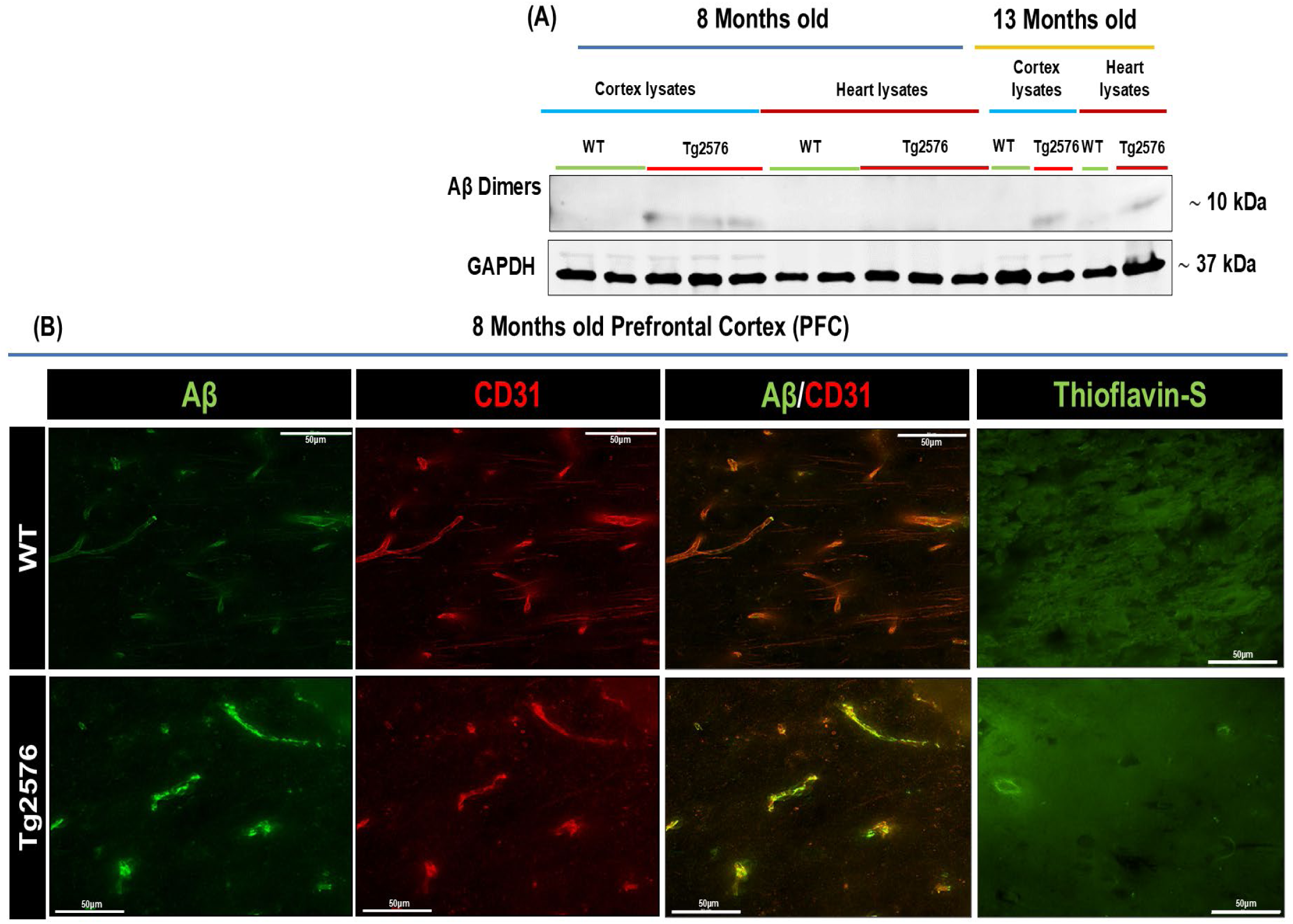
Amyloid-β deposits in vessels of 8 months-old Tg2576 prefrontal cortex (PFC), but not in heart tissue. (**A**) Representative immunoblots show Aβ dimers in total cardiac lysates from WT and Tg2576 mice. Tg2576 cerebral cortex lysate was used as an internal positive control. GAPDH levels were used as a loading control. (**B**) Representative digital images (scale bar 50μm) showing vessels (stained with CD31, in red) and Aβ deposits (in green). Right panels: representative digital images (right panels; scale bar 50μm) showing β-sheet conformation (fibrillar) amyloid marked with Thioflavin-S (in green) in the prefrontal cortex (PFC), in 8-months old WT and Tg2576 mice.

**Supplemental Figure 6.**
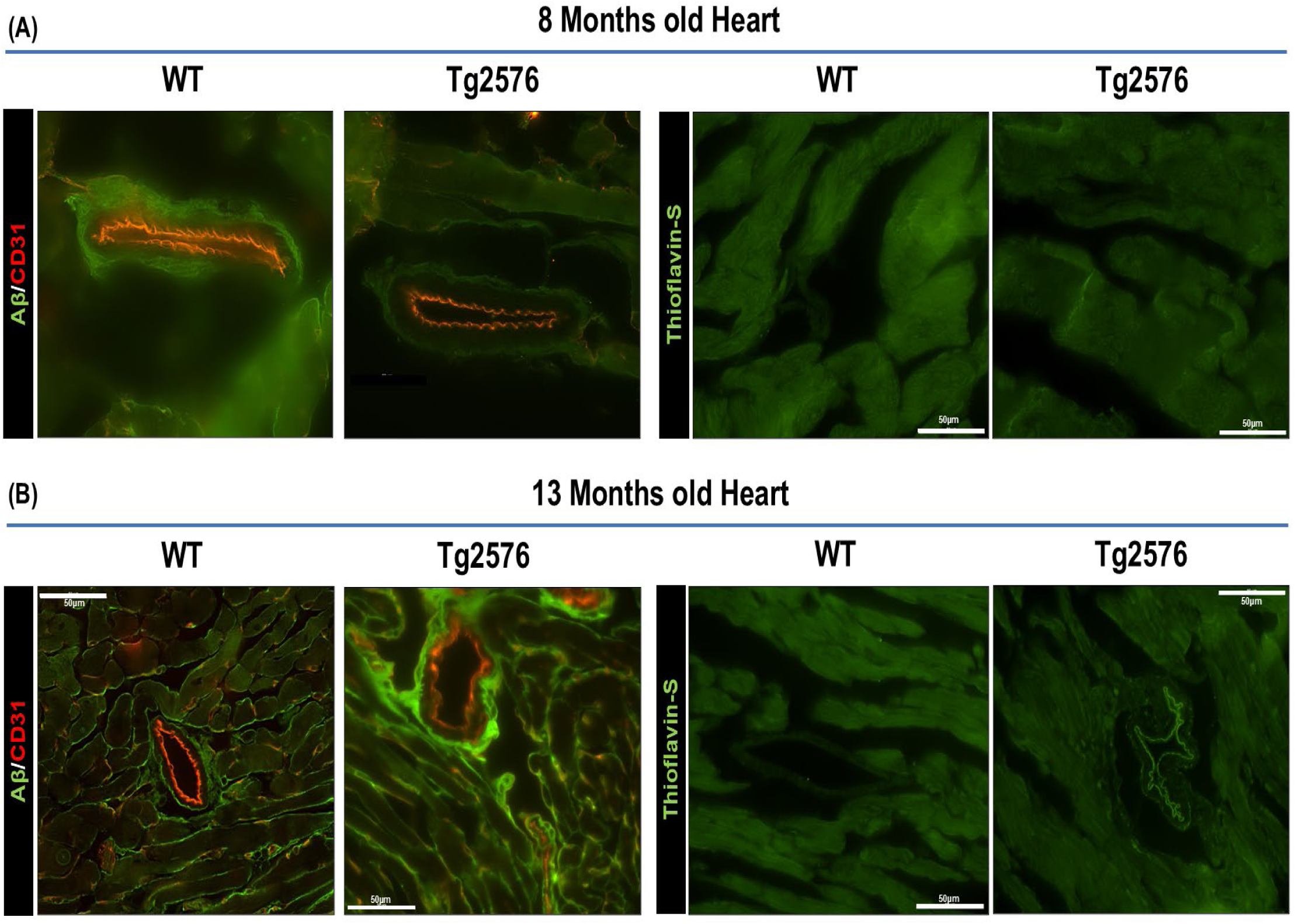
AD pathology affects the cardiac vascular network in 13-month-old Tg2576 mice. (**A**-**B**) Representative digital images (left panels, scale bar 50μm) showing vessels (stained with CD31, in red) and Aβ deposits (in green), and representative digital images (right panels; scale bar 50μm) showing β-sheet conformation of amyloid marked with Thioflavin-S (in green) in cardiac sections from WT and Tg2576 mice.

**Supplemental Figure 7.**
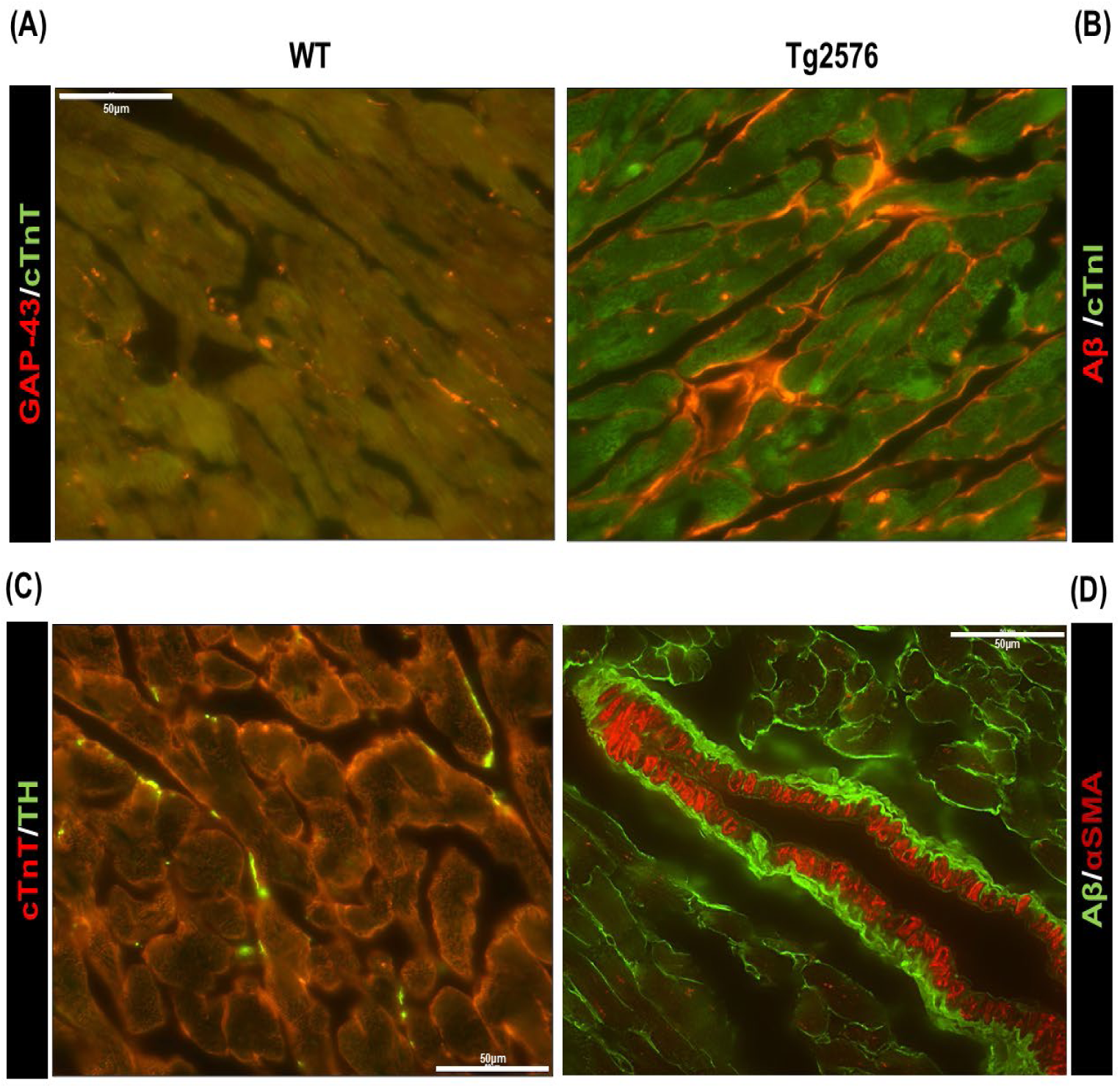
Representative digital images (scale bar 50μm) of cardiac sections from 13-month-old WT and Tg2576 mice stained with. (**A**) anti-neuronal regeneration marker (GAP-43, in red) and cardiac Troponin T (cTnT, in green). (**B**) Aβ (in red) and cardiac Troponin I (cTnI, in green). (**C**) anti-tyrosine-hydroxylase (TH, in green) and cardiac Troponin T (cTnT, in red). (**D**) Aβ (in green) and α-smooth muscle actin (α-SMA, in red).

**Supplemental Figure 8.**
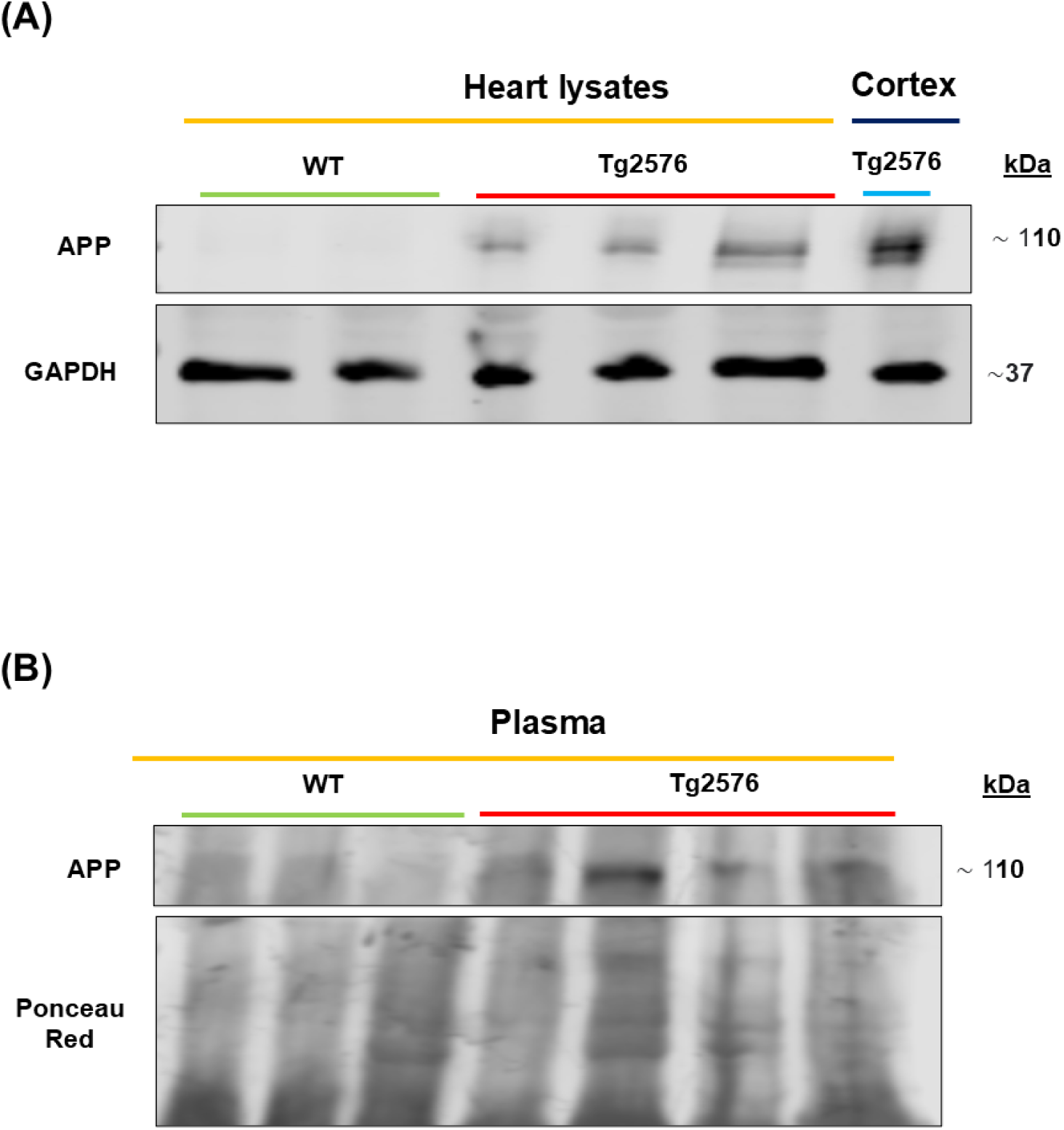
The Amyloid precursor protein (APP) is present in plasma and heart tissue of the 13-month-old Tg2576-AD model. (**A**-**B**) Representative immunoblots showing protein levels of APP in total cardiac lysates (upper panel) and plasma APP levels (lower panel) in 13-month-old WT and Tg2576 mice. Tg2576 cortex lysate was used as an internal positive control. GAPDH levels or ponceau red were used as respective loading controls.

**Supplemental Figure 9.**
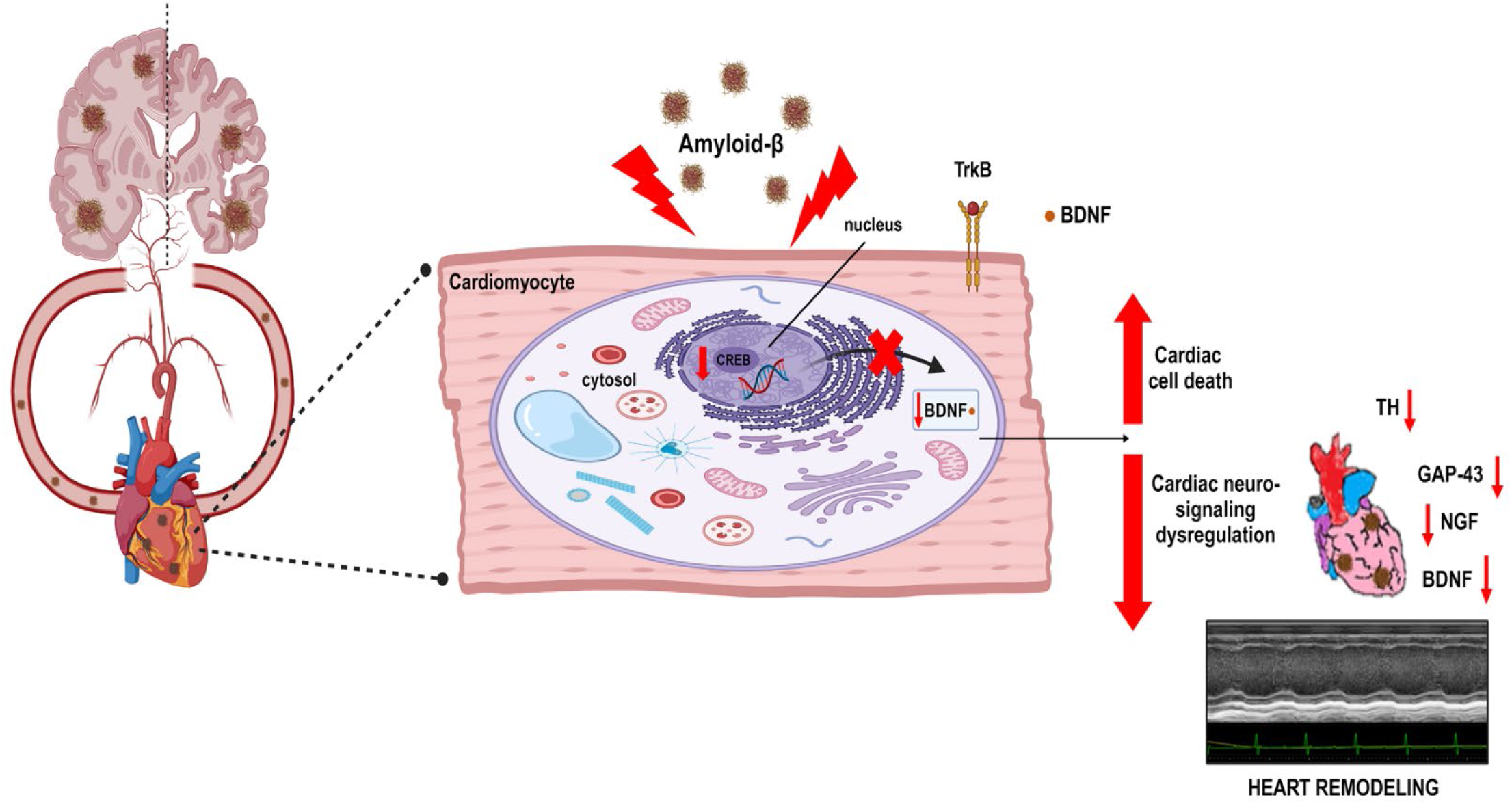
Conceptual scheme of the study. Amyloid-β aggregates, present in the heart parenchyma of AD mouse models and AD human hearts, induce adverse cardiac remodeling and myocardial denervation through cardiac TrkB/CREB/BDNF neurotrophic signaling axis dysregulation, ultimately resulting in impaired cardiac physiology.

